# Impaired neuromodulator crosstalk delays vigilance-dependent astroglia Ca^2+^ activation in mouse models of Alzheimer’s disease

**DOI:** 10.1101/2022.06.27.497832

**Authors:** Eunice Y. Lim, Angelica Salinas, Liang Ye, Yongjie Yang, Martin Paukert

**Affiliations:** Department of Cellular and Integrative Physiology, University of Texas Health Science Center at San Antonio, San Antonio, TX, United States of America; Center for Biomedical Neuroscience, University of Texas Health Science Center at San Antonio, San Antonio, TX, United States of America; Department of Neuroscience, Tufts University, Boston, MA, United States of America

## Abstract

Degeneration in neuronal nuclei producing the neuromodulators acetylcholine and norepinephrine is a hallmark of Alzheimer’s disease (AD). Therapeutic interventions that increase acetylcholine in brain ameliorate AD symptoms in human patients, and augmenting norepinephrine restores cognitive function in mouse models of AD as well as Down Syndrome, the most frequent cause of early onset AD. A prominent cellular target of noradrenergic and potentially cholinergic signaling during states of heightened vigilance are astroglia and recent studies indicate that astroglia Ca^2+^ dynamics in awake mice contribute to optimal cognitive performance. Here we tested the hypothesis that vigilance-dependent Ca^2+^ signaling in mouse primary visual cortex astrocytes is altered in mouse models of AD and provide mechanistic insight into upstream neuromodulator signaling that shapes astrocyte Ca^2+^ dynamics in healthy and AD conditions. In two mouse models of AD (APPswe/PSEN1dE9 and *App^NL-F^ KI*), we consistently observed delayed and less coordinated astrocyte Ca^2+^ elevations in response to locomotion, a well-controlled behavioral paradigm triggering widespread Ca^2+^ activation in astroglia throughout the brain. Combining pharmacological and genetic manipulations, we found that noradrenergic signaling to astrocytes was facilitated by cholinergic signaling, but this neuromodulator crosstalk was impaired in *App^NL-F^* mice. Pharmacological facilitation of norepinephrine release rescued delayed and less coordinated astrocyte Ca^2+^ activation in *App^NL-F^* mice and suggests that astrocytes preserve a functional reserve that can be recruited even during late-stage disease. Our findings of delayed and less coordinated astroglia Ca^2+^ activation predict impaired noradrenergic signaling and may contribute to the cognitive decline in AD.

## Introduction

Alzheimer’s disease (AD), the most common cause of dementia, is accompanied by prominent anatomical and neurochemical alterations in neuromodulator systems. Human necropsy brain samples revealing an inverse correlation of severity of dementia or senile plaque load with expression level of choline acetyl transferase (ChAT), the rate-limiting enzyme of acetylcholine synthesis, inspired the cholinergic hypothesis of geriatric memory dysfunction (Bartus, Dean, Beer, & Lippa, 1982; Bergmann, Tomlinson, Blessed, Gibson, & Perry, 1978; PERRY, PERRY, BLESSED, & TOMLINSON, 1978). Many mouse models of AD replicate reduced ChAT activity in hippocampus and cortex and some undergo degeneration of basal forebrain cholinergic neurons (Shekari & Fahnestock, 2021). Enhancing levels of acetylcholine in the brain by inhibiting its degradation by acetylcholinesterase has remained the mainstay of AD treatment strategies once dementia has manifested (Bartus, 2000; Mufson, Counts, Perez, & Ginsberg, 2008). In addition to the cholinergic system, neurons in locus coeruleus, the primary source of norepinephrine in the brain, undergo degeneration in AD patients. While locus coeruleus neuron loss is uniformly diffuse in elderly people (Marcyniuk, Mann, & Yates, 1989), it is confined to the dorsal portion of locus coeruleus in Alzheimer’s and Down syndrome patients with Down syndrome representing the most frequent cause for early-onset AD (Marcyniuk, Mann, & Yates, 1986; Marcyniuk, Mann, Yates, & Ravindra, 1988). Degeneration of noradrenergic neurons is not only a consequence of AD but also influences its disease course. In mouse models of AD and Down syndrome, chemical or genetic impairment of locus coeruleus function exacerbated hallmarks of AD: Aβ plaque deposition, neurofibrillary tangles formation, inflammation, synaptic dysfunction, neurodegeneration or cognitive decline (Hammerschmidt et al., 2013; Heneka et al., 2002, 2006; Jardanhazi-Kurutz et al., 2011; Lockrow et al., 2011). In contrast, supporting noradrenergic signaling by administering the norepinephrine prodrug L-threo-3,4-dihydroxyphenylserine (L-DOPS) restored contextual memory function in a Down syndrome mouse model (Salehi et al., 2009) and improved cognitive and biochemical parameters in an AD mouse model (Kalinin et al., 2012). The cellular mechanisms of how noradrenergic signaling benefits cognitive performance in AD-related mouse models are not completely understood but recently glial cells (Braun, Madrigal, & Feinstein, 2014), and in particular astrocyte Ca^2+^ dynamics, have become a focus of interest as potential mediators (Verkhratsky, 2019; Zorec, Parpura, & Verkhratsky, 2018).

In response to locomotion, mouse astroglia undergo global Ca^2+^ elevations that extend across the whole cell including soma and processes, across the astroglia population (Dombeck, Khabbaz, Collman, Adelman, & Tank, 2007; Nimmerjahn, Mukamel, & Schnitzer, 2009), and occur simultaneously in regions of the brain as different as visual cortex and cerebellum (Gray, Ye, Ye, & Paukert, 2021; Paukert et al., 2014). Locomotion excites noradrenergic as well as cholinergic nerve terminals (Reimer et al., 2016) and triggers release of the respective neuromodulators (Fu et al., 2014; Paukert et al., 2014; Polack, Friedman, & Golshani, 2013); and electrical stimulation of locus coeruleus or nucleus basalis of Meynert revealed that both norepinephrine as well as acetylcholine can trigger robust Ca^2+^ elevations in cortical astrocytes (Bekar, He, & Nedergaard, 2008; Chen et al., 2012; Takata et al., 2011). Yet, a predominant dependence on α_1_-adrenergic signaling has consistently been revealed by pharmacological analysis of astroglia Ca^2+^ elevations induced by locomotion or other means of provoking states of heightened vigilance (Ding et al., 2013; Paukert et al., 2014; Srinivasan et al., 2015). More specifically, gene deletion of the α_1A_-adrenergic receptor subtype from cerebellar Bergmann glia revealed its requirement for locomotion-induced Ca^2+^ elevations (Ye et al., 2020).

Consistent with the possibility of direct noradrenergic action on cortical astrocytes, most of the heightened vigilance-dependent Ca^2+^ elevations depend on G_q_ protein-coupled receptor signaling (Nagai et al., 2021) and subsequent release of Ca^2+^ from the endoplasmic reticulum via inositol trisphosphate receptor type 2 (IP_3_R2) (Srinivasan et al., 2015). This is important because manipulations of astrocyte G_q_ protein- or G_i/o_ protein-coupled receptor signaling, which both lead to IP_3_R2-dependent Ca^2+^ elevations (Durkee et al., 2019), or gene deletion of IP_3_R2 have been found to impact select cognitive functions. Effects on remote memory function have been discovered following gene deletion of IP_3_R2 (Pinto-Duarte, Roberts, Ouyang, & Sejnowski, 2019), activation of G_i/o_ protein-coupled receptors in hippocampal astrocytes (Kol et al., 2020), or transient activation of G_q_ protein-coupled receptor in frontal cortex astrocytes (Iwai et al., 2021). In striatal astrocytes, activation of G_i/o_ protein-coupled receptors leads to general hyperactivity (Nagai et al., 2019), and inhibition of G_q_ protein-coupled receptor signaling impairs spatial memory (Nagai et al., 2021). Therefore, precisely balanced astroglia Ca^2+^ signaling is required for normal cognitive performance.

We hypothesized that cortical astrocyte Ca^2+^ dynamics associated with a behavioral state of heightened vigilance might be affected by altered neuromodulator signaling in mouse models of AD. We found that locomotion-induced visual cortex astrocyte Ca^2+^ elevations were consistently delayed and less coordinated in two mouse models of AD. A considerable portion of this phenomenon could be accounted for by impaired cholinergic facilitation of norepinephrine release which could be partially restored by pharmacological enhancement of noradrenergic terminal excitability. Our findings suggest that disturbed vigilance-dependent astroglia Ca^2+^ dynamics due to impaired crosstalk between cholinergic and noradrenergic signaling may contribute to cognitive decline in AD.

## Results

### Delayed and less coordinated vigilance-dependent V1 astrocyte Ca^2+^ elevations in APP/PS1 mice

In order to inform our understanding of potential deficits underlying the Alzheimer’s phenotype in astrocytes, we observed astrocyte Ca^2+^ dynamics in mouse primary visual cortex (V1) during long periods of locomotion in a model of Alzheimer’s disease and dissected the upstream signaling mechanisms that give rise to these Ca^2+^ dynamics. Throughout monitoring of V1 astrocyte Ca^2+^ dynamics, mice were kept in darkness to minimize complications in interpreting data originating from modulation of signals by processing of local sensory information (Gray et al., 2021; Slezak et al., 2019). We used *in vivo* two-photon (2P) microscopy with a previously reported setup of head-fixation beneath the objective and positioning on a linear motorized treadmill (Paukert et al., 2014) and observed Ca^2+^ dynamics in astrocytes of left V1 during 2-minute bouts of enforced locomotion (Figure 1A). While astroglia Ca^2+^ transients exhibit similar kinetics during voluntary or enforced locomotion (Paukert et al., 2014), the advantage of using enforced locomotion to analyze astrocyte Ca^2+^ signaling is that it removes the uncertainty in determining the start of arousal, which with voluntary locomotion, may occur anywhere from when the animal intends to move to actual movement. Thus, this allows precise quantification of the kinetics of Ca^2+^ transients relative to stimulus onset.

**Figure 1.**
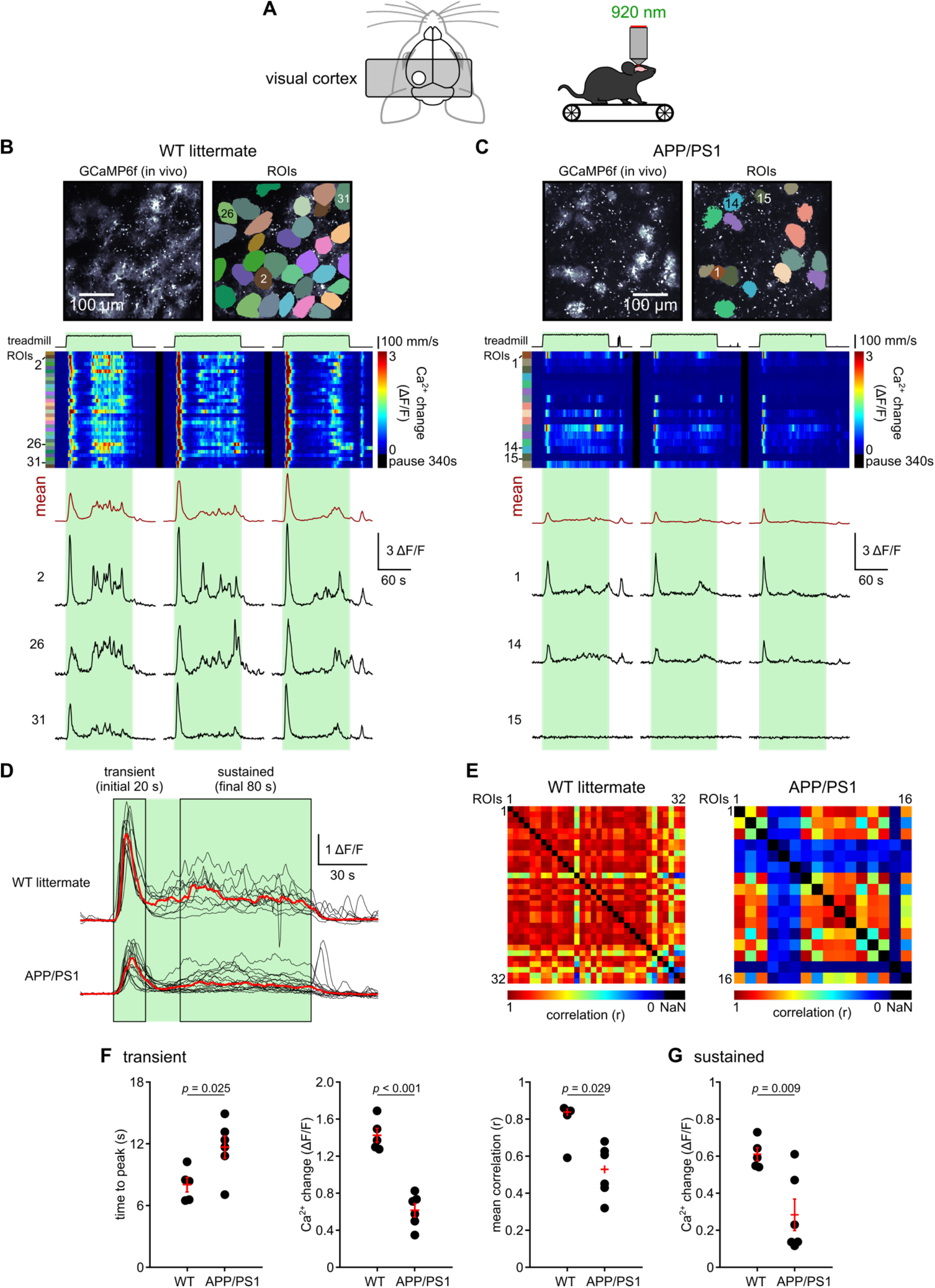
Mouse model for Alzheimer’s disease exhibits slower and reduced vigilance-dependent V1 astrocyte Ca^2+^ dynamics. (**A**) **Left**, schematic showing placement of steel headplate and chronic cranial window above V1 of left hemisphere. **Right**, schematic of *in vivo* 2P Ca^2+^ imaging in awake, head-restrained mouse on a motorized linear treadmill. (**B**) **Top left**, image of *in vivo* GCaMP6f in V1 astrocytes in a representative APP/PS1 wildtype littermate expressing *Slc1a3-CreER^T^;Ai95*, tangential optical section of ~60 μm beneath pial surface. **Top right**, regions of interest (ROIs). Lower, astrocyte Ca^2+^ signals during 3 bouts of enforced locomotion from representative mouse. **Top to bottom**, ‘treadmill’, speed of treadmill; pseudocolor plot of Ca^2+^ responses in all ROIs indicated at top right; ‘mean’ (dark red), average of all ROIs’ Ca^2+^ response traces; ‘2’, ‘26’, ‘31’, example traces of ROIs 2, 26, and 31; green bars, enforced locomotion. (**C**) Same as **B** in a representative APP/PS1 *;Slc1a3-CreER^T^;Ai95* mouse. (**D**) Overlay of aligned Ca^2+^ fluorescence traces from APP/PS1 wildtype littermates (*n* = 11 traces each representing the average Ca^2+^ response of all ROIs in a FOV, 5 mice) and APP/PS1 mice (*n* = 16 traces, 6 mice). Red trace is median of individual traces. (**E**) **Left**, mean correlation coefficient across all ROIs during initial 20 seconds of locomotion, or transient phase in same representative APP/PS1 wildtype littermate as in **B**. **Right**, same as left in same representative APP/PS1 mouse as in **C**. (**F**) Population data. APP/PS1 wildtype littermates, *n* = 5 mice; APP/PS1, *n* = 6 mice. **Left**, time to peak; unpaired, two-tailed Student’s *t*-test (*t(9)* = −2.693, *p* = 0.025). **Center**, mean Ca^2+^ change during transient phase; unpaired, two-tailed Student’s *t*-test (*t*(9) = 7.706, *p* < 0.001). **Right**, mean correlation coefficient during transient phase; Kruskal Wallis test (*p* = 0.029). (**G**) Mean Ca^2+^ change during last 80 seconds of locomotion, or sustained phase; unpaired, two-tailed Student’s *t*-test (*t*(9) = 3.306, *p* = 0.009). (**F-G**) If all data follow Gaussian distribution red symbols indicate mean ± SEM, if not red symbols represent median.

APP/PS1 transgenic mice (17-22 months) were used as a model of familial Alzheimer’s disease. APP/PS1 mice overexpress in CNS neurons 3 times the endogenous level of APP the chimeric mouse/human amyloid-β precursor protein with Swedish mutations K595N and M596L (Mo/HuAPP695swe) and a mutant human presenilin 1 carrying the exon-9-deleted variant PSEN1dE9; this results in beta-amyloid deposits by 6-7 months of age (Jankowsky et al., 2004). To monitor cytosolic Ca^2+^ dynamics, APP/PS1 mice were bred to co-express astrocyte-specific CreER^T^ (Paukert et al., 2014) and floxed-stop GCaMP6f (Ai95) (Madisen et al., 2015), resulting in *APP/PS1;Slc1a3-CreER^T^;Ai95* mice. Age-matched wildtype littermates lacking APP/PS1 were used as controls. Astrocyte Ca^2+^ dynamics were analyzed at the single-cell level by assigning regions of interest (ROIs) encompassing the entire cell including the soma and processes (Figure 1B and C).

Following a synchronized population-wide global astrocyte Ca^2+^ elevation that has been described above (analyzed during the first 20 s of each locomotion trial), as locomotion continued, we observed sustained irregularly sized and timed Ca^2+^ elevations lasting the duration of locomotion (analyzed during last 80 s of each locomotion trial; Figure 1B-D). Averaging of responses of all astrocytes in a field-of-view (FOV) revealed a steady-state population Ca^2+^ response of various amplitudes during continuous locomotion (Figure 1D). In APP/PS1 mice, heightened vigilance-dependent astrocyte Ca^2+^ dynamics were delayed and less coordinated (Figure 1D-F), and the transient as well as the sustained responses were reduced in mean amplitude (Figure 1F and G) compared to wildtype littermates. Occasionally, non-responsive astrocytes could be found in APP/PS1 mice (Figure 1C; e.g. ROI ‘15’). The marked deficits of astrocyte Ca^2+^ signaling in this model of AD may be due to impaired upstream signaling or reduced responsiveness of astrocytes or both.

### In normal young adult mice, vigilance-dependent V1 astrocyte Ca^2+^ elevations are delayed by inhibition of cholinergic signaling

ChAT activity in cortex of APP/PS1 mice is already reduced in mice at 10-16 months of age, which is earlier than the mice used in our study (Perez, Dar, Ikonomovic, DeKosky, & Mufson, 2007). Therefore, we considered impaired upstream signaling to account for disrupted vigilance-dependent V1 astrocyte Ca^2+^ dynamics in this mouse model of AD and first tested whether inhibition of acetylcholine receptors induces similar changes in astrocyte Ca^2+^ dynamics in young adult wildtype mice. While hippocampal astrocytes respond to acetylcholine via nicotinic receptors (Pabst et al., 2016; Papouin, Dunphy, Tolman, Dineley, & Haydon, 2017), Ca^2+^ responses of cortical astrocytes to electrical stimulation of nucleus basalis of Meynert are dependent on muscarinic receptors (Chen et al., 2012; Takata et al., 2011). Therefore, we explored the effect of the muscarinic inhibitor scopolamine on V1 astrocyte Ca^2+^ dynamics during locomotion. In 2-7 month-old *Adra1a^+/+^;Slc1a3-CreER^T^;Ai95* mice, similar Ca^2+^ dynamics were observed as in the aged wildtype littermates of APP/PS1 mice shown in Figure 1, demonstrating that the 2-component astrocyte response is maintained throughout young adulthood and old age in mice. Scopolamine had different effects on the transient and sustained components. Blocking muscarinic receptors through scopolamine (3 mg/kg, i.p.) reduced the rate of Ca^2+^ rise during the transient component and the mean amplitude of the sustained component (Figure 2A, C and D). There was only a trend towards less synchronized transient responses (Figure 2B and C) and, consistent with findings from vigilance-dependent cerebellar Bergmann glia Ca^2+^ elevations (Paukert et al., 2014), inhibition of muscarinic receptors did not significantly reduce the mean amplitude of the transient response (Figure 2C). Interestingly, this impairment induced by acute inhibition of muscarinic receptors resembled baseline responses in APP/PS1 mice, though less severely. Together, this data indicates that during vigilance, acetylcholine indirectly or directly participates in the astrocyte Ca^2+^ response by accelerating its rise and enhancing its steady-state response during continuous alertness. It remains to be determined whether the muscarinic receptors are activated directly on astrocytes, as has previously been proposed (Sugihara, Chen, & Sur, 2016; Takata et al., 2011), or on upstream intermediary neurons.

**Figure 2.**
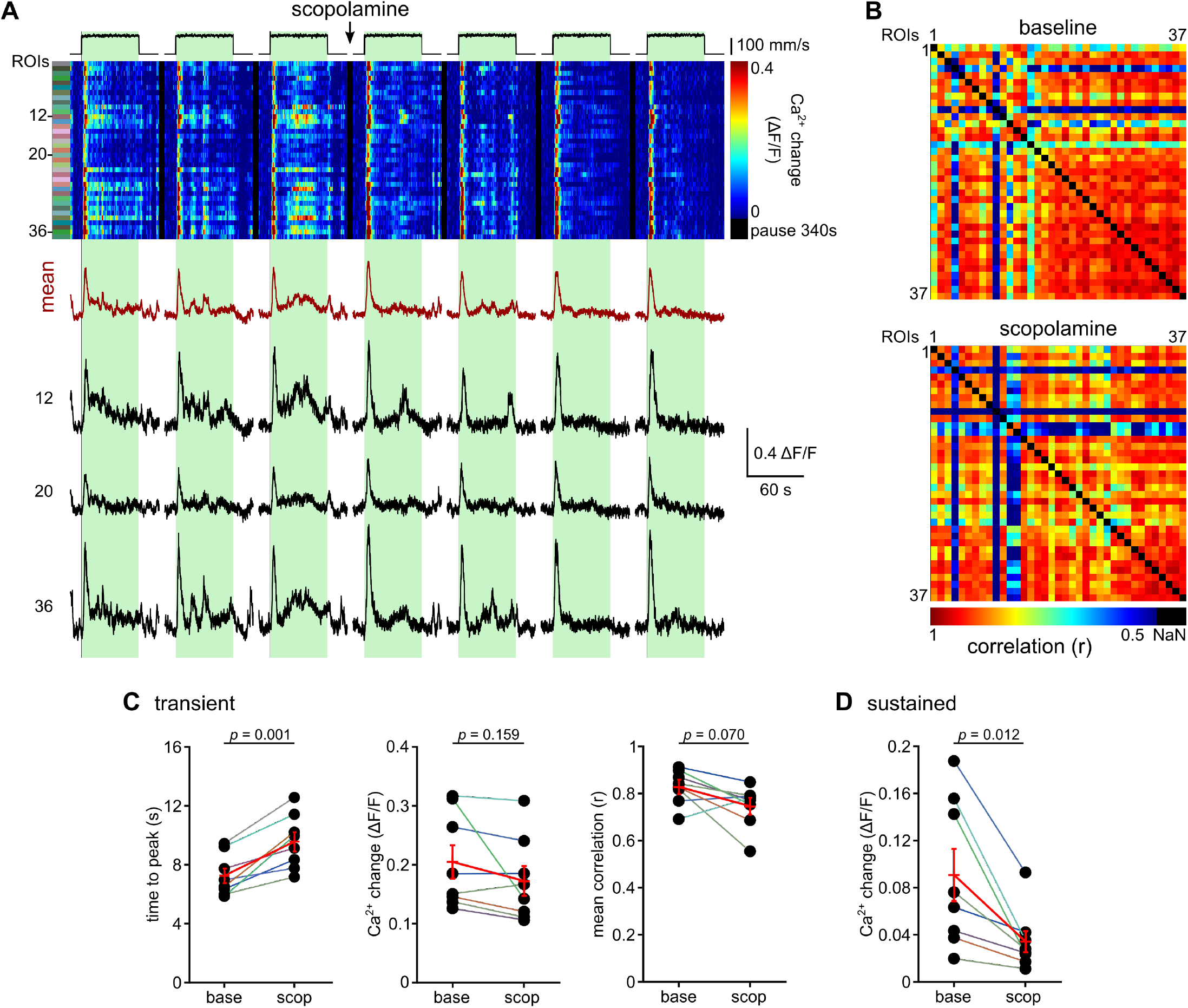
Inhibition of muscarinic receptors causes slower initial Ca^2+^ transient and reduced dynamics during sustained portion of locomotion. (**A**) Pseudocolor plot of Ca^2+^ responses in representative mouse showing timecourse of experiment with administration of muscarinic antagonist scopolamine (3 mg/kg, i.p.); ‘mean’ (dark red), average of all ROIs’ Ca^2+^ response traces; ‘12’, ‘20’, ‘36’, example traces of respective ROIs; green bars, enforced locomotion. (**B**) Mean correlation coefficient across ROIs during transient locomotion phase of before (top) and after scopolamine injection (bottom) in the representative mouse. (**C**) Population data: **left**, time to peak; paired, two-tailed Student’s *t*-test (*t*(7) = −5.373, *p* = 0.001); *n* = 8. **Center**, mean Ca^2+^ change; paired, two-tailed Student’s *t*-test (*t*(7) = 1.575, *p* = 0.159). **Right**, mean correlation coefficient; paired, two-tailed Student’s *t*-test (*t*(7) = 2.137, *p* = 0.070). (**D**) Population data: mean Ca^2+^ change during sustained phase; paired, two-tailed Student’s *t*-test (*t*(7) = 3.363, *p* = 0.012). (**C-D**) All data follow Gaussian distribution, and red symbols indicate mean ± SEM.

### α_1A_-adrenergic receptors are required for the majority of vigilance-dependent V1 astrocyte Ca^2+^ elevations and the residual signal is not sensitive to inhibition of cholinergic signaling

Arousal-associated cortical astrocyte Ca^2+^ signaling depends on α_1_-adrenergic receptors (Ding et al., 2013; Srinivasan et al., 2015) and in cerebellar Bergmann glia on α_1A_-adrenergic receptors (Ye et al., 2020). Having just established the involvement of acetylcholine, we aimed to dissect the relative contribution of noradrenergic and cholinergic signaling to vigilance-dependent V1 astrocyte Ca^2+^ dynamics and to determine whether both simultaneously and directly activate astrocytes or whether they are involved in a signaling hierarchy. To do this, we investigated α_1A_-adrenergic receptor global knockout mice (Rokosh & Simpson, 2002) comparing *Adra1a;Slc1a3-CreER^T^;Ai95* mice that were mutant, heterozygous or wildtype for *Adra1a* and tested the effect of various pharmacological drugs in these transgenic mice. Under baseline conditions, heterozygous mice behaved like wildtype, but complete loss of α_1A_-adrenergic receptor function resulted in Ca^2+^ responses that took almost twice as long to reach the peak and were considerably less coordinated. Furthermore, the mean amplitude of the transient response was reduced to a one third residual response while the steady-state response to continuous locomotion was almost completely abolished (Figure 3A-C). The latter finding explicitly indicates that the astrocyte steady-state Ca^2+^ elevations in this continuous locomotion paradigm are mediated by norepinephrine.

**Figure 3.**
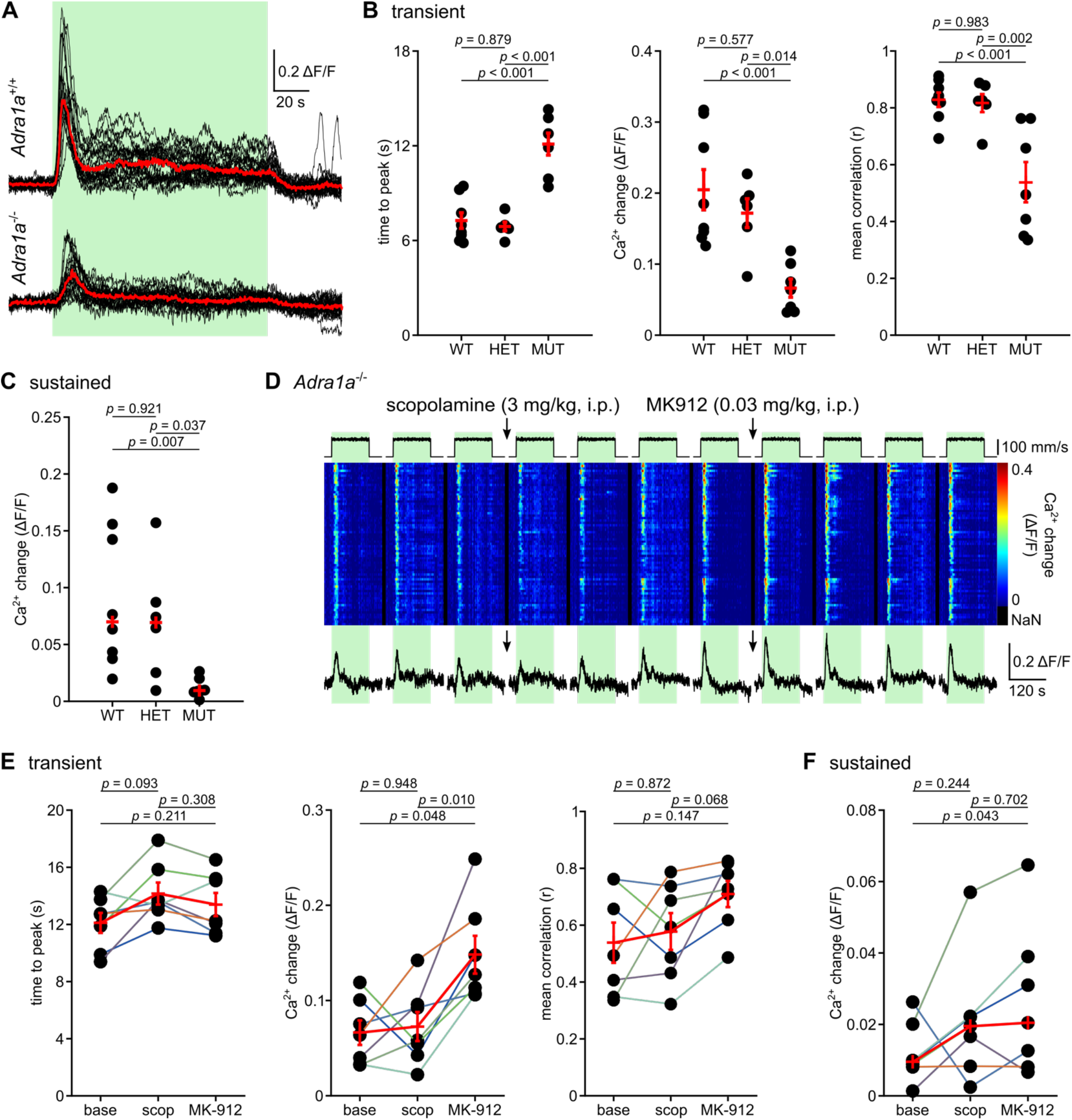
Genetic knockdown of α_1A_-adrenergic receptors dramatically reduced speed and amplitude of vigilance-dependent V1 astrocyte Ca^2+^ dynamics, and the residual Ca^2+^ signals in *Adra1a^-/-^* mice were unaffected by muscarinic inhibition. (**A**) Overlay of aligned Ca^2+^ fluorescence traces in *Adra1a^+/+^* mice (*n* = 22 traces, 8 mice) and *Adra1a^-/-^* mice (*n* = 21 traces, 7 mice). Red traces represent median of respective individual traces. (**B**) Population data: *Adra1a^+/+^, n* = 8 mice; *Adra1a^+/-^, n* = 6 mice; *Adra1a^-/-^, n* = 7 mice. **Left,** time to peak; one-way ANOVA followed by Tukey-Kramer correction (F(2,18) = 28.691, *p* < 0.001). **Center,** mean Ca^2+^ change; one-way ANOVA followed by Tukey-Kramer correction (F(2,18) = 10.555, *p* < 0.001). **Right**, mean correlation coefficient; one-way ANOVA followed by Tukey-Kramer correction (F(2,18) = 12.330, *p* < 0.001). (**C**) Population data: mean Ca^2+^ change during sustained phase; Kruskal Wallis test followed by Tukey-Kramer correction (*p* = 0.005). (**D**) Pseudocolor plot of Ca^2+^ responses in representative *Adra1a^-/-^* mouse showing timecourse of experiment with administration of muscarinic antagonist scopolamine (3 mg/kg, i.p) and α_2_-adrenergic receptor antagonist MK-912 (0.03 mg/kg, i.p.). Trace represents mean of all ROIs’ Ca^2+^ responses. (**E**) Population data: **Left**, time to peak; repeated measures ANOVA followed by Tukey-Kramer correction (F(2,12) = 4.947, *p* = 0.027). **Center**, mean Ca^2+^ change; repeated measures ANOVA followed by Tukey-Kramer correction (F(2,12) = 8.903, *p* = 0.004). **Right**, mean correlation coefficient; repeated measures ANOVA followed by Tukey-Kramer correction (F(2,12) = 3.370, *p* = 0.069). (**F**) Population data: mean Ca^2+^ change during sustained phase; Friedman test followed by Tukey-Kramer correction (χ^2^(2) = 6, *p* = 0.050). (**B-C**, **E-F**) If all data follow Gaussian distribution red symbols indicate mean ± SEM, if not red symbols represent median.

To test whether the residual vigilance-dependent V1 astrocyte Ca^2+^ response in α_1A_-adrenergic receptor knockout mice represented a cholinergic response, we administered scopolamine. However, besides a trend towards further delay in Ca^2+^ responses, scopolamine had no effect on residual astrocyte Ca^2+^ dynamics in α_1A_-adrenergic receptor knockout mice (Figure 3D-F). Instead, the α_2_-adrenergic receptor antagonist MK-912, which antagonizes autoinhibitory presynaptic receptors and facilitates norepinephrine release (Ye et al., 2020), enhanced the mean transient response amplitude and caused a trend towards increasing coordination (Figure 3D-E), suggesting that the residual astrocyte Ca^2+^ response is mediated by non-A subtype α1-adrenergic receptors. Together, these data support a model where, upon transition to a more vigilant state in mice, acetylcholine facilitates norepinephrine release, which via activation of predominantly α_1A_-adrenergic receptors, leads to a transient Ca^2+^ elevation that is followed by a still norepinephrine-driven steady-state Ca^2+^ elevation for the duration of heightened vigilance. This ‘cholinergic facilitation model’ would predict that acetylcholine-mediated facilitation of vigilance-dependent norepinephrine release is reflected in noradrenergic terminal Ca^2+^ activity in V1.

### Vigilance-dependent noradrenergic terminal Ca^2+^ elevations are impaired by inhibition of cholinergic signaling

To test this prediction, Ca^2+^ dynamics in noradrenergic terminals of V1 were observed during locomotion. Membrane-tethered GCaMP6f was expressed in noradrenergic neurons by crossing *Dbh-Cre* (Gerfen, Paletzki, & Heintz, 2013) and *Lck-GCaMP6f*^flox^ (Srinivasan et al., 2016) mice. Membrane-tethering is required in these transgenic mice to detect noradrenergic terminal Ca^2+^ dynamics with GCaMP6f (Ye et al., 2020). Since individual nerve terminals cannot be recognized, we used an unbiased checkerboard arrangement of 50 μm x 50 μm ROIs (Figure 4A; (Gray et al., 2021; Ye et al., 2020)). Noradrenergic terminals exhibited a similar Ca^2+^ response as astrocytes with two components - a fast, synchronized transient component followed by sustained activity (Figure 4B and C). Further similar to astrocytes, Ca^2+^ transients could be triggered by enforced or voluntary locomotion (Figure 4B, arrowheads). The reduced sustained activity indicates that the astrocyte Ca^2+^ response attenuation to the steady-state level is not primarily limited by astrocyte responsiveness. Consistent with the predictions from the ‘cholinergic facilitation model’, scopolamine slowed down vigilance-dependent V1 noradrenergic terminal Ca^2+^ rise and reduced mean amplitude and spatial correlation of the population transient response (Figure 4D-G). Unexpectedly, scopolamine did not reduce the mean amplitude of the steady-state Ca^2+^ dynamics during continuous locomotion (Figure 4G); possible explanations will be presented in the discussion.

**Figure 4.**
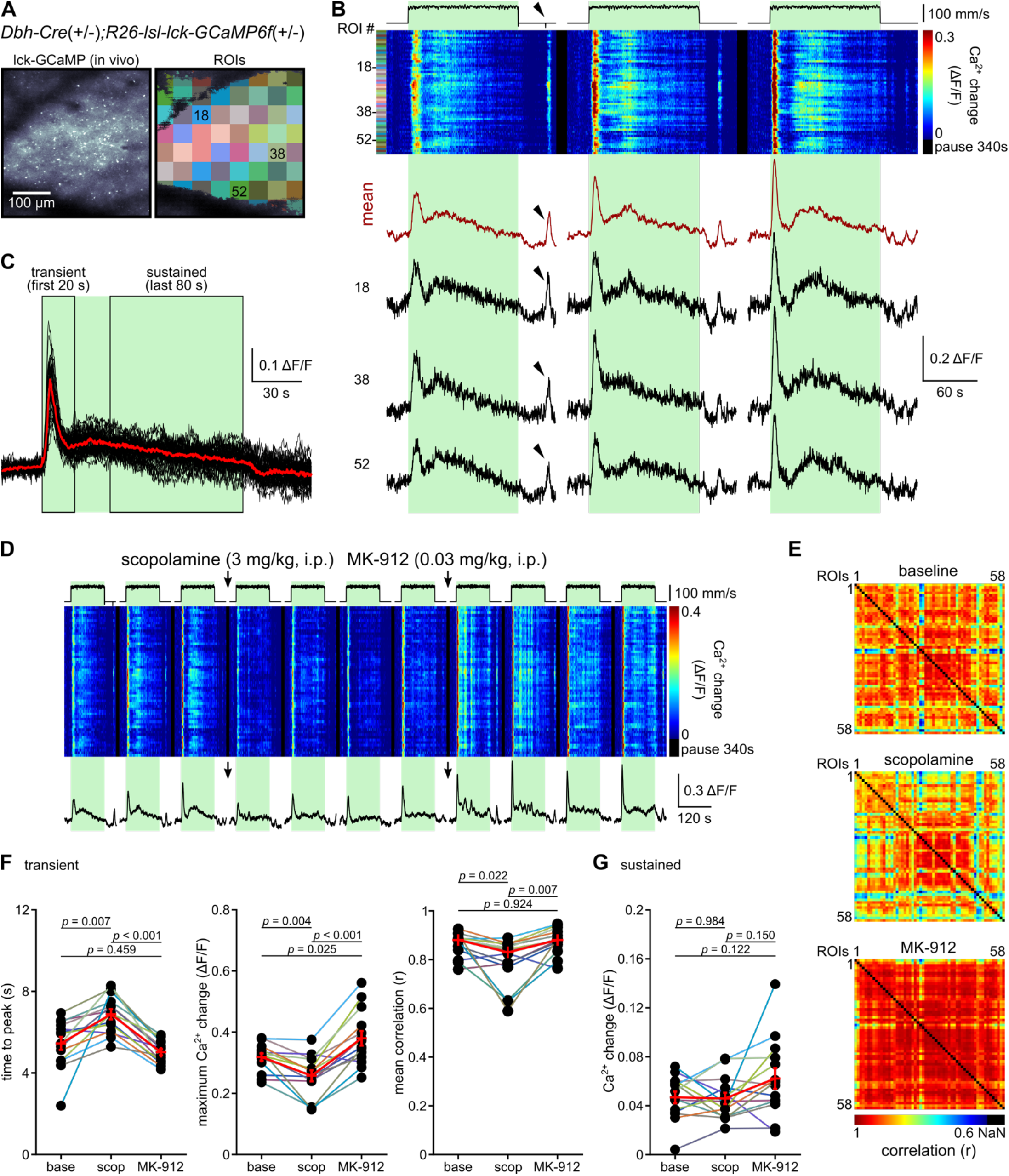
Vigilance-dependent Ca^2+^ signals in noradrenergic terminals were slower and reduced during muscarinic receptor inhibition. (**A**) **Left**, image of *in vivo* Lck-GCaMP6f in noradrenergic terminals of V1 in a representative *Dbh-Cre;Lck-GCaMP6f^flox^* mouse, tangential optical section of ~60 μm beneath pial surface. **Right**, regions of interest (ROIs). (**B**) Noradrenergic terminal Ca^2+^ signals during 3 bouts of enforced locomotion from representative mouse. **Top to bottom**: speed of treadmill; pseudocolor plot of Ca^2+^ responses in all ROIs indicated in **A**; ‘mean’ (dark red), average of all ROIs’ Ca^2+^ response traces; ‘18’, ‘38’, ‘52’, example traces of respective ROIs; green bars, enforced locomotion; black arrows indicates a bout of voluntary movement. (**C**) Overlay of aligned Ca^2+^ fluorescence traces (*n* = 40 traces, 14 mice). Red trace is median of individual traces. (**D**) Pseudocolor plot and mean trace of representative mouse showing timecourse of experiment with administration of muscarinic antagonist scopolamine (3 mg/kg, i.p.) and α_2_-adrenergic receptor antagonist MK-912 (0.03 mg/kg, i.p.). (**E**) Maps of correlation coefficients across ROIs during repetitions before scopolamine injection (‘baseline’), after scopolamine and before MK-912 injections (‘scopolamine’), and following MK-912 injection (‘MK-912’). (**F**) **Left**, time to peak; repeated measures ANOVA followed by Tukey-Kramer correction (F(2,26) = 17.948, *p* < 0.001). **Center**, mean Ca^2+^ change; repeated measures ANOVA followed by Tukey-Kramer correction (F(2,26) = 23.646, *p* < 0.001). **Right**, mean correlation coefficient; Friedman test followed by Tukey-Kramer correction (χ^2^(2) = 10.857, *p* = 0.004). (**G**) Mean Ca^2+^ change during sustained phase; repeated measures ANOVA followed by Tukey-Kramer correction (F(2,26) = 3.483, *p* = 0.046). (**F-G**) If all data follow Gaussian distribution red symbols indicate mean ± SEM, if not red symbols represent median.

In summary, the finding that noradrenergic terminal Ca^2+^ activity is inhibited by scopolamine together with the considerable reduction of nearly all aspects of Ca^2+^ activity in α_1A_-adrenergic receptor knockout mice and the resistance of the residual responses in those mice to scopolamine (Figure 3), support the model of cholinergic facilitation of noradrenergic signaling to V1 astrocytes. The similarity between the impairment caused by acute muscarinic receptor inhibition and the deficit under baseline conditions in the APP/PS1 mouse model of AD raised the possibility that impaired cholinergic facilitation is a mechanistic contributor to disrupted astrocyte Ca^2+^ dynamics in AD mouse models. To test this possibility and further explore how consistent changes to vigilance-dependent cortical astrocyte Ca^2+^ dynamics are among mouse models of AD, we investigated a mouse model of slow-progressing AD.

### Delay of vigilance-dependent V1 astrocyte Ca^2+^ activation and loss of its cholinergic facilitation in *App^NL-F^* mice

The *App^NL-F^* model consists of knock-in expression (replacing endogenous mouse APP) of humanized murine APP with Swedish KM670/671NL and Beyreuther/Iberian I716F mutations and maintains endogenous APP expression level. It has been characterized as a slow-progressing AD model that better mimics human AD progression (Saito et al., 2014). Aged *App^NL-F/NL-F^* mice and age-matched C57BL/6J wildtype control mice (16-22 month-old) were tested based on availability. Mice were injected with AAV-PHP.B-gfaABC1D-GCaMP_3_ virus, which contains the blood-brain-barrier penetration facilitating capsid PHP.B (Deverman et al., 2016) and has GCaMP_3_ expression controlled by the astrocyte-specific promoter gfaABC1D (Lee, Messing, Su, & Brenner, 2008), into the retro-orbital venous sinus. A previous study demonstrates the compatibility of studying astrocyte Ca^2+^ dynamics with either GCaMP6f or GCaMP_3_ (Ye, Haroon, Salinas, & Paukert, 2017). Time-intense pharmacological experiments testing the effects of scopolamine and MK-912 were modified, and astrocyte Ca^2+^ responses were observed during locomotion bouts lasting 5 seconds instead of 2 minutes.

Both the rate of rise of vigilance-dependent V1 astrocyte Ca^2+^ elevation as well as the correlation of responses among astrocytes were attenuated in *App^NL-F/NL-F^* mice but not the mean amplitude (Figure 5A-E). This response profile was similar but weaker than that of APP/PS1 mice (Figure 1), which was expected in a slow-progressing model of AD, and mimicked the profile seen in wildtype mice when muscarinic receptor signaling was acutely inhibited (Figure 2). The *App^NL-F/NL-F^* phenotypes of slower time to peak and reduced inter-astrocyte correlation were further analyzed for the pharmacological effects of inhibiting muscarinic receptors via scopolamine and boosting norepinephrine release through MK-912. Aged wildtype mice recapitulated the scopolamine-mediated deceleration of Ca^2+^ rise and the trend towards reduced correlation among individual astrocytes’ responses (Figure 5F) seen in adult wildtype mice (Figure 2C). Strikingly, any effect of scopolamine was lost in *App^NL-F/NL-F^* mice. This result even sustained analysis of single astrocytes with its increased statistical power (Figure 5G). This means that cholinergic facilitation of norepinephrine release may have been disrupted at this disease stage.

**Figure 5.**
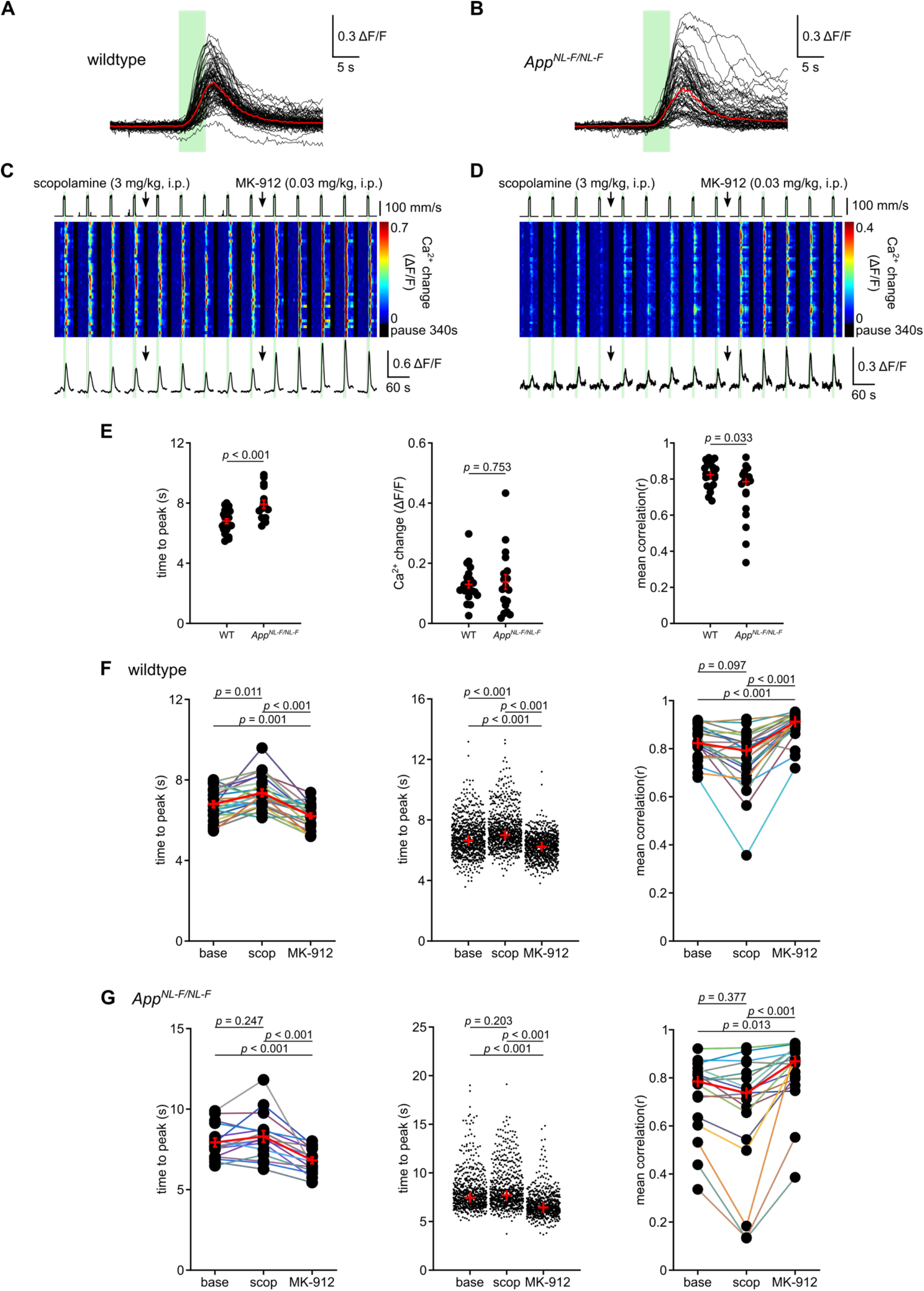
*App^NL-F/NL-F^* knock-in model of Alzheimer’s disease exhibits slower vigilance-dependent V1 astrocyte Ca^2+^ signals compared to age-matched wildtype mice, and the slower responses are not effected by muscarinic inhibition. (**A**) Overlay of aligned Ca^2+^ fluorescent traces in aged wildtype mice (*n* = 79 traces, 8 mice). (**B**) Same as in A for aged *App^NL-F/NL-F^* mice (*n* = 65 traces, 7 mice). (**C**) Pseudocolor plot of Ca^2+^ responses showing timecourse of experiment with administration of muscarinic antagonist scopolamine (3 mg/kg, i.p.) and α_2_-adrenergic receptor antagonist MK-912 (0.03 mg/kg, i.p.); trace, average of all ROIs’ Ca^2+^ response traces; green bars, enforced locomotion. (**D**) Same as **C** for a representative aged *App^NL-F/NL-F^* mice. (**E**) **Left**, population data: time to peak; aged wildtype, *n* = 22 FOVs, 8 mice; aged *App^NL-F/NL-F^, n* = 16 FOVs, 7 mice; unpaired, two-tailed Student’s *t*-test (*t*(36) = −3.860, *p* < 0.001). **Center**, mean Ca^2+^ change; aged wildtype, *n* = 23 FOVs, 8 mice; aged *App^NL-F/NL-F^, n* = 18 FOVs, 7 mice; unpaired, two-tailed Student’s *t*-test (*t*(39) = −0.317, *p* = 0.753). **Right**, mean correlation coefficient; aged wildtype, *n* = 23 FOVs, 8 mice; aged *App^NL-F/NL-F^, n* = 18 FOVs, 7 mice; Kruskal Wallis test (*p* = 0.033). (**F**) **Left**, population data: time to peak in aged wildtype mice; *n* = 22 FOVs, 8 mice; repeated measures ANOVA followed by Tukey-Kramer correction (F(2,42) = 24.901, *p* < 0.001). **Center**, population data: time to peak in individual ROIs of aged wildtype mice (*n* = 824 ROIs, 8 mice); Friedman test followed by Tukey-Kramer correction (χ^2^(2) = 511.756, *p* < 0.001). **Right**, population data: mean correlation coefficient across ROIs in aged wildtype mice; *n* = 23 FOVs, 8 mice; Friedman test and Tukey-Kramer correction (χ^2^(2) = 35.826, *p* < 0.001). (**G**) **Left**, population data: time to peak in aged *App^NL-F/NL-F^* mice; *n* = 16 FOVs, 7 mice; repeated measures ANOVA followed by Tukey-Kramer correction (F(2,30) = 18.339, *p* < 0.001). **Center**, population data: time to peak in individual ROIs of aged *App^NL-F/NL-F^* mice (*n* = 678 ROIs, 7 mice); Friedman test and Tukey-Kramer correction (χ^2^(2) = 535.369, *p* < 0.001). **Right**, population data: mean correlation coefficient across ROIs in aged *App^NL-F/NL-F^* mice; *n* = 18 FOVs, 7 mice; Friedman test and Tukey-Kramer correction (χ^2^(2) = 18.111, *p* < 0.001). (**E-G**) If all data follow Gaussian distribution red symbols indicate mean ± SEM, if not red symbols represent median.

The apparent loss of sensitivity to scopolamine indicates a deficit somewhere in the signaling hierarchy, either at the level of cholinergic facilitation or on noradrenergic terminal excitability or on astrocyte responsivity. To determine if the deficit lay in cholinergic facilitation, MK-912 was used to pharmacologically intervene immediately downstream by enhancing noradrenergic terminal excitability. This resulted in improved time to peak and correlation among individual astrocytes’ responses in *App^NL-F/NL-F^* mice as well as in wildtype mice (Figure 5F and G). The improvement was sufficient to restore wildtype baseline levels while it did not reach the effect of MK-912 on wildtype animals (Supplementary Figure 1). The observation that MK-912 could restore wildtype baseline levels while *App^NL-F/NL-F^* mice were insensitive to scopolamine indicates that cholinergic facilitation was impaired.

## Discussion

In this study we have investigated heightened vigilance-dependent V1 astrocyte Ca^2+^ dynamics in two mouse models of AD, a model of accelerated pathology (APP/PS1) and a milder knock-in mouse model (*App^NL-F/NL-F^*). Consistently in both models, we found that at onset of locomotion, the astrocyte Ca^2+^ elevation rose more slowly and reached the peak amplitude later. In addition, the correlation among Ca^2+^ responses of individual astrocytes was reduced, indicating less coordinated population activity. Combining gene deletion with *in vivo* pharmacology experiments, we made a number of key observations that allowed us to build a model of crosstalk between cholinergic and noradrenergic signaling to explain a considerable portion of this phenotype in AD mouse models. (1) Acute pharmacological inhibition of muscarinic receptors in wildtype mice mimicked the AD phenotype - deceleration of astrocyte Ca^2+^ elevations and the reduced coordination of population activity. (2) α_1A_-adrenergic receptor knockout mice revealed that the majority (approximately two thirds) of the initial transient and almost all of the sustained phase of astrocyte responses required activation of this receptor. Importantly, the residual responses following α_1A_-adrenergic receptor knockout were not significantly affected by inhibition of muscarinic receptors indicating that direct activation of muscarinic receptors on cortical astrocytes does not significantly contribute to vigilance-dependent whole astrocyte Ca^2+^ elevations. Instead, the residual responses were responsive to pharmacological facilitation of norepinephrine release and exhibited enhanced mean amplitude and coordination, suggesting that non-A subtype α1-adrenergic receptors account for the residual response. (3) The pharmacological effects of acute inhibition of muscarinic receptors on V1 noradrenergic terminal Ca^2+^ response mirrored those of astrocytes, demonstrating that noradrenergic neurons are regulated by muscarinic receptor activation in this behavioral context. (4) The *App^NL-F/NL-F^* phenotype regarding vigilance-dependent astrocyte Ca^2+^ dynamics was characterized by lack of inhibition by muscarinic receptor antagonism, suggesting that further inhibition of this signaling pathway was occluded by an inherent deficit in cholinergic signaling in these mice. (5) For both mouse models of AD at progressed stage, acute pharmacological facilitation of vigilance-triggered norepinephrine release could partially rescue normal astrocyte Ca^2+^ dynamics. Together these findings support a model of impaired cholinergic facilitation of vigilance-dependent norepinephrine release leading to slower cortical astrocyte Ca^2+^ activation in mouse models of AD. The data further suggest that noradrenergic signaling to astrocytes possesses a functional reserve that can be pharmacologically recruited.

The rate of Ca^2+^ rise, measured as *time* from onset of locomotion *to peak* of the Ca^2+^ transient, has emerged as a sensitive and reliable parameter to assess acute or chronic changes of vigilance-dependent excitation of noradrenergic terminals and Ca^2+^ activation of astroglia. Acute exposure to ethanol in mice causes longer time to peak of noradrenergic terminal as well as Bergmann glia vigilance-dependent Ca^2+^ elevations by inhibiting excitation of noradrenergic neurons and their cerebellar terminals; both observations can be reversed by the α_2_-adrenergic receptor antagonist MK-912 (Ye et al., 2020). Accordingly, visual stimulation-triggered short-term potentiation of V1 noradrenergic terminals is accompanied by shorter time to peak of vigilance-dependent Ca^2+^ elevations in noradrenergic terminals and astrocytes (Gray et al., 2021).

Our study provides a behavioral context in which cholinergic innervation of noradrenergic neurons may be particularly important. A mechanism describing how acetylcholine facilitates norepinephrine release may be that acetylcholine activates muscarinic receptors expressed on locus coeruleus noradrenergic neurons and increases their excitability. Evidence for this is found in acute brain slices of rat locus coeruleus where muscarine reduces inward rectifier potassium conductance of noradrenergic neurons and thereby increases their excitability (Shen & North, 1992). Additionally, the same study also showed that muscarine caused an inward current around resting membrane potential. Therefore, under intense activation of cholinergic neurons, such as electrical stimulation of nucleus basalis of Meynert, cholinergic signaling may be sufficient to trigger norepinephrine release and account for at least a portion of the astrocyte Ca^2+^ elevations in anesthetized mice (Chen et al., 2012; Takata et al., 2011). Since nicotinic acetylcholine receptors can contribute to excitation in rat locus coeruleus neurons (Egan & North, 1986), the extent of cholinergic facilitation of noradrenergic signaling to astrocytes revealed by inhibition of muscarinic receptors (Figure 2) may still represent an underestimation. Our study shows that direct activation of muscarinic receptors on astrocytes by acetylcholine is unlikely to significantly contribute to these responses since inhibition of muscarinic receptors had no effect in α_1A_-adrenergic receptor knockout mice (Figure 3). However, it must be stated that direct activation of astrocyte muscarinic receptors may contribute to astrocyte microdomain Ca^2+^ activity.

Remarkably, our ‘cholinergic facilitation model’ of norepinephrine release may help explain some phenomena that have been observed at the microdomain level, which are triggered by diverse mechanisms (Agarwal et al., 2017; Lim, Ye, & Paukert, 2021). When muscarinic receptors are inhibited, the average amplitude of whisker stimulation-enhanced microdomain Ca^2+^ activity in barrel cortex astrocytes is not affected, but their onset latency is significantly prolonged (Stobart et al., 2018). This nuanced effect of altering kinetics and not amplitude resembles the delayed global responses induced by scopolamine in our study and suggests that cholinergic facilitation of norepinephrine release may not only give rise to vigilance-dependent global Ca^2+^ responses but also a portion of microdomain events. While some can be triggered by sensory stimulation, the rest occur spontaneously; plasma membrane ion channels appear to account for faster events (Stobart et al., 2018) but a large proportion also depends on IP_3_R2 (Agarwal et al., 2017; Srinivasan et al., 2015). Consistent with the spontaneous nature, microdomain Ca^2+^ events dominate astroglia Ca^2+^ dynamics during anesthesia (Nimmerjahn et al., 2009; Thrane et al., 2012). Astrocyte microdomain Ca^2+^ events occur in close proximity to mitochondria (Jackson & Robinson, 2015; Stephen et al., 2015), require transient opening of the mitochondrial permeability transition pore, and can be potentiated by reactive oxygen species and were enhanced in a mouse model carrying a mutation in superoxide dismutase 1 (SOD1) associated with a familial form of amyotrophic lateral sclerosis (Agarwal et al., 2017). An interesting similarity was observed in anesthetized APP/PS1 where cortical astrocyte microdomain Ca^2+^ events were more frequent and synchronous (Kuchibhotla, Lattarulo, Hyman, & Bacskai, 2009), a feature that was consistent across subpopulations from three other mouse models of AD at early disease stages and associated with unstable vascular tone (Takano, Han, Deane, Zlokovic, & Nedergaard, 2007). Pharmacological analysis of the spontaneous Ca^2+^ activity in astrocytes of anesthetized APP/PS1 mice revealed that paracrine purinergic signaling via P2Y1 receptors preferentially near amyloid plaques accounts for most of the hyperactivity (Delekate et al., 2014). The considerable contrast between astrocyte Ca^2+^ hyperactivity under anesthesia and delayed, less coordinated astrocyte Ca^2+^ activation in awake behaving mice induced by heightened vigilance emphasizes the importance of considering the underlying mechanisms of distinct forms of astrocyte Ca^2+^ dynamics associated with different behavioral and environmental context.

Our finding that both astrocyte as well as noradrenergic terminal Ca^2+^ elevations declined to a reduced steady-state level following an initial transient response indicates that astrocyte responsiveness and intracellular Ca^2+^ release are not the primary limitation of sustained Ca^2+^ activation. It was then surprising that inhibition of muscarinic receptors led to a robust inhibition of the V1 astrocyte sustained Ca^2+^ response (Figure 2D), but the same pharmacological treatment had no effect on the sustained Ca^2+^ response of noradrenergic terminals (Figure 4G). One explanation could be that the sustained astrocyte Ca^2+^ response is mediated by acetylcholine and not norepinephrine; however, the α_1A_-adrenergic receptor knockout data indicate that the vast majority of the sustained astrocyte response depends on noradrenergic signaling (Figure 3A). Another explanation could be that noradrenergic terminal Ca^2+^ signals monitored with membrane-anchored GCaMP6f better represents presynaptic Ca^2+^ available for vesicular release rather than Ca^2+^ available for endocytosis and vesicle pool recovery. It is expected that vesicle pool recovery becomes more relevant during sustained locomotion to maintain norepinephrine release. Norepinephrine release from sympathetic neurons depends on N-type Ca^2+^ channels (Hirning et al., 1988), and in contrast to L-type Ca^2+^ channels, N-type Ca^2+^ channels are tethered to the active zone and releasable vesicles via RIM proteins (Kaeser et al., 2011). Interestingly, for a *Drosophila* glutamatergic synapse, it has been demonstrated that Ca^2+^ dynamics controlled by P/Q- or N-type Ca^2+^ channel homologs are involved in exocytosis and separated by plasma membrane Ca^2+^ ATPase (PMCA) from Ca^2+^ dynamics controlled by L-type Ca^2+^ channel homologs, which have been found to be important for endocytosis (Krick et al., 2021). If this presynaptic functional organization represented a general principle also for the mammalian neurotransmitter release machinery, it might be interesting in future studies to tether GCaMPs selectively to protein complexes containing N-type versus L-type Ca^2+^ channels to monitor vesicular release and recovery separately.

We found that enhancing norepinephrine release through the α_2_-adrenergic autoreceptor inhibitor MK-912 was sufficient to restore vigilance-dependent V1 astrocyte Ca^2+^ signaling in *App^NL-F/NL-F^* mice to wildtype baseline levels but not to the full potential that was reached by MK-912 in wildtype mice (Supplementary Figure 1). This indicates that impairments other than cholinergic facilitation are also involved. Human tissue from AD patients revealed 50 - 60% neuronal loss in locus coeruleus (German et al., 1992; Matthews et al., 2002), preferentially in its dorsal portion (Marcyniuk et al., 1986), which would predict a lower density of noradrenergic terminals in projection areas. Consistently, the density of noradrenergic axon terminals in cerebral cortex is reduced in APP/PS1 and *App^NL-G-F/NL-G-F^* mice (Cao, Fisher, Rodriguez, Yu, & Dong, n.d.; Sakakibara et al., 2021). It is then conceivable that reduced density of norepinephrine release sites in V1, which would limit the amount of achievable norepinephrine release even if noradrenergic terminals were maximally excited, accounts for MK-912 potentiating vigilance-dependent astrocyte Ca^2+^ responses in wildtype mice further than in *App^NL-F/NL-F^* mice.

In conclusion, we have obtained functional evidence for cholinergic facilitation of heightened vigilance-dependent noradrenergic signaling to V1 astrocytes. In light of the rapidly emerging role of astroglia Ca^2+^ dynamics in cognitive function (Dallérac & Rouach, 2016; Kofuji & Araque, 2021; Santello, Toni, & Volterra, 2019), the impairment of this behavioral state-dependent neuromodulator crosstalk at late stage of two mouse models of AD (Figure 6) suggests that impaired vigilance-dependent astrocyte Ca^2+^ activation may contribute to cognitive decline in AD. Our findings that acute inhibition of α_2_-adrenergic receptors was able to restore a considerable portion of astrocyte Ca^2+^ activation suggests that there is a functional reserve for norepinephrine release as well as astrocyte responsiveness that can be recruited. α_2_-adrenergic receptor antagonists are possibly effective since they do not only facilitate norepinephrine release by inhibiting autoreceptors but they may also facilitate acetylcholine release (Williams & Reiner, 1993). Therefore, α_2_-adrenergic receptor antagonism may offer a pharmacological strategy to support the heightened vigilance-dependent neuromodulatory crosstalk and signaling to noradrenergic targets including astrocytes in AD.

**Figure 6.**
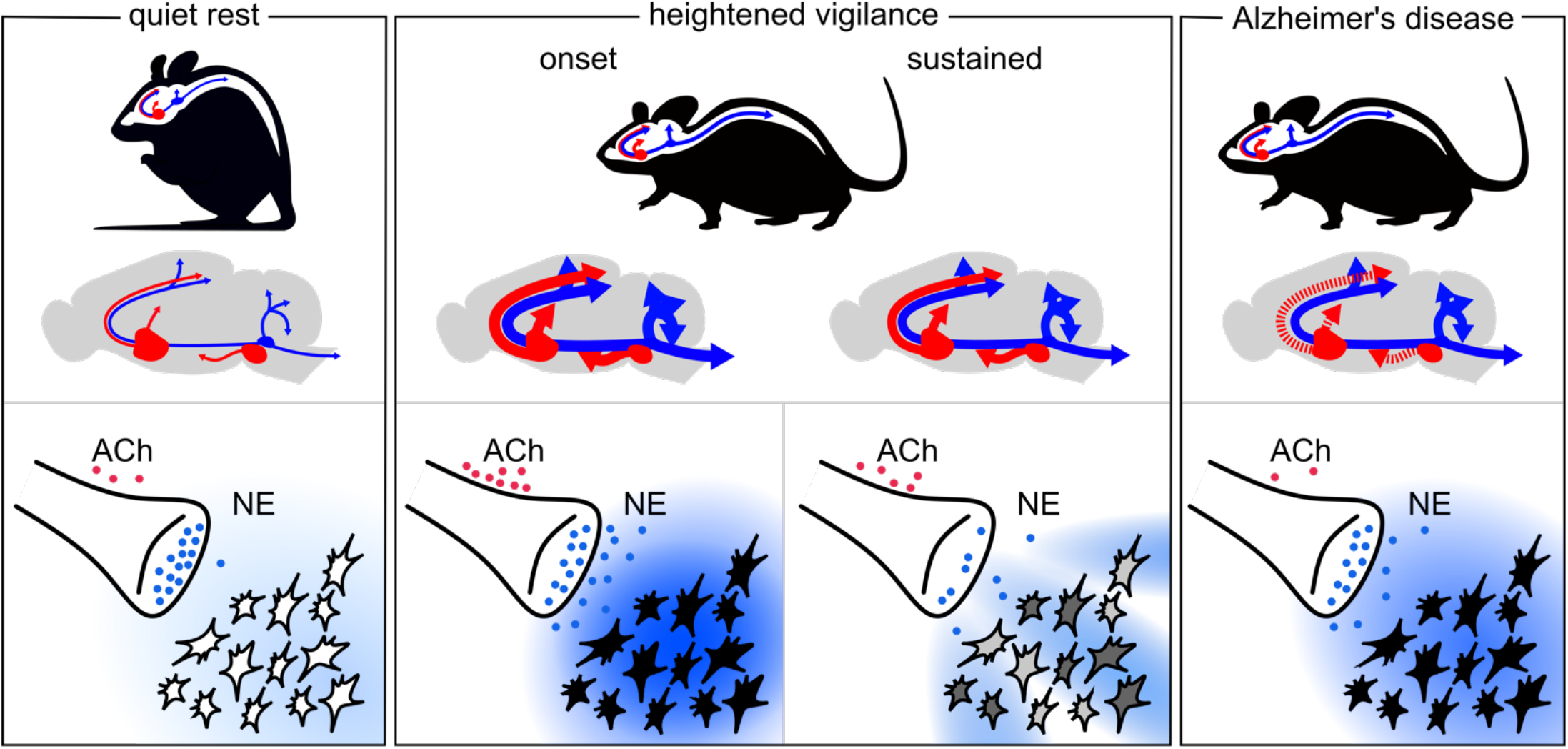
Schematic of our proposed ‘cholinergic facilitation model’. Cholinergic system is shown in red, noradrenergic system is shown in blue. Acetylcholine is symbolized as red dots and norepinephrine as blue dots or clouds. Gray scale represents different level of astrocyte Ca^2+^ activation.

## Acknowledgments

The authors would like to thank Dr. Takaomi Saido for permission to use the *App^NL-F^* mouse line, Drs. Viviana Gradinaru and Ben Deverman for sharing the AAV-GFAP-mKate2.5f and AAV-PHP.B plasmids before these were publicly available, Raehum Paik from the UTHSCSA Mouse Genome Engineering and Transgenic Facility for support with the AAV-PHP.B-gfaABC1D-GCaMP_3_ construct and Priscilla M. Barba-Escobedo and John Cavaretta for providing expert support in genotyping and animal husbandry. Eunice Y. Lim was supported by a T32NS082145 fellowship. This work was supported by R01MH113780 and The Robert J. Kleberg, Jr. and Helen C. Kleberg Foundation (Martin Paukert).

## References

Agarwal, A., Wu, P. H., Hughes, E. G., Fukaya, M., Tischfield, M. A., Langseth, A. J., … Bergles, D. E. (2017). Transient Opening of the Mitochondrial Permeability Transition Pore Induces Microdomain Calcium Transients in Astrocyte Processes. Neuron, 93(3), 587–605.e7. https://doi.org/10.1016/J.NEURON.2016.12.034/ATTACHMENT/7E42C560-E2D8-402B-BE94-068438DF30FF/MMC10.PDF

Bartus, R. T. (2000). On neurodegenerative diseases, models, and treatment strategies: lessons learned and lessons forgotten a generation following the cholinergic hypothesis. Experimental Neurology, 163(2), 495–529. https://doi.org/10.1006/EXNR.2000.7397

Bartus, R. T., Dean, R. L., Beer, B., & Lippa, A. S. (1982). The cholinergic hypothesis of geriatric memory dysfunction. Science (New York, N.Y.), 217(4558), 408–417. https://doi.org/10.1126/SCIENCE.7046051

Bekar, L. K., He, W., & Nedergaard, M. (2008). Locus coeruleus alpha-adrenergic-mediated activation of cortical astrocytes in vivo. Cerebral Cortex (New York, N.Y.: 1991), 18(12), 2789–2795. https://doi.org/10.1093/CERCOR/BHN040

Bergmann, K., Tomlinson, B. E., Blessed, G., Gibson, P. H., & Perry, R. H. (1978). Correlation of cholinergic abnormalities with senile plaques and mental test scores in senile dementia. British Medical Journal, 2(6150), 1457. https://doi.org/10.1136/BMJ.2.6150.1457

Braun, D., Madrigal, J., & Feinstein, D. (2014). Noradrenergic regulation of glial activation: molecular mechanisms and therapeutic implications. Current Neuropharmacology, 12(4), 342–352. https://doi.org/10.2174/1570159X12666140828220938

Cao, S., Fisher, D. W., Rodriguez, G., Yu, T., & Dong, H. (n.d.). Comparisons of neuroinflammation, microglial activation, and degeneration of the locus coeruleus-norepinephrine system in APP/PS1 and aging mice. https://doi.org/10.1186/s12974-020-02054-2

Chen, N., Sugihara, H., Sharma, J., Perea, G., Petravicz, J., Le, C., & Sur, M. (2012). Nucleus basalis-enabled stimulus-specific plasticity in the visual cortex is mediated by astrocytes. Proceedings of the National Academy of Sciences of the United States of America, 109(41). https://doi.org/10.1073/PNAS.1206557109/-/DCSUPPLEMENTAL/PNAS.201206557SI.PDF

Dallérac, G., & Rouach, N. (2016). Astrocytes as new targets to improve cognitive functions. Progress in Neurobiology, 144, 48–67. https://doi.org/10.1016/J.PNEUROBIO.2016.01.003

Delekate, A., Füchtemeier, M., Schumacher, T., Ulbrich, C., Foddis, M., & Petzold, G. C. (2014). Metabotropic P2Y1 receptor signalling mediates astrocytic hyperactivity in vivo in an Alzheimer’s disease mouse model. Nature Communications 2014 5:1, 5(1), 1–14. https://doi.org/10.1038/ncomms6422

Deverman, B. E., Pravdo, P. L., Simpson, B. P., Kumar, S. R., Chan, K. Y., Banerjee, A., … Gradinaru, V. (2016). Cre-dependent selection yields AAV variants for widespread gene transfer to the adult brain. Nature Biotechnology 2015 34:2, 34(2), 204–209. https://doi.org/10.1038/nbt.3440

Ding, F., O’Donnell, J., Thrane, A. S., Zeppenfeld, D., Kang, H., Xie, L., … Nedergaard, M. (2013). α1-Adrenergic receptors mediate coordinated Ca2+ signaling of cortical astrocytes in awake, behaving mice. Cell Calcium, 54(6), 387–394. https://doi.org/10.1016/J.CECA.2013.09.001

Dombeck, D. A., Khabbaz, A. N., Collman, F., Adelman, T. L., & Tank, D. W. (2007). Imaging large-scale neural activity with cellular resolution in awake, mobile mice. Neuron, 56(1), 43–57. https://doi.org/10.1016/J.NEURON.2007.08.003

Durkee, C. A., Covelo, A., Lines, J., Kofuji, P., Aguilar, J., & Araque, A. (2019). Gi/o protein-coupled receptors inhibit neurons but activate astrocytes and stimulate gliotransmission. Glia, 67(6), 1076–1093. https://doi.org/10.1002/GLIA.23589

Egan, T. M., & North, R. A. (1986). Actions of acetylcholine and nicotine on rat locus coeruleus neurons in vitro. Neuroscience, 19(2), 565–571. https://doi.org/10.1016/0306-4522(86)90281-2

Fu, Y., Tucciarone, J. M., Espinosa, J. S., Sheng, N., Darcy, D. P., Nicoll, R. A., … Stryker, M. P. (2014). A cortical circuit for gain control by behavioral state. Cell, 156(6), 1139–1152. https://doi.org/10.1016/J.CELL.2014.01.050

Gerfen, C. R., Paletzki, R., & Heintz, N. (2013). GENSAT BAC cre-recombinase driver lines to study the functional organization of cerebral cortical and basal ganglia circuits. Neuron, 80(6), 1368–1383. https://doi.org/10.1016/J.NEURON.2013.10.016

German, D. C., Manaye, K. F., White, C. L., Woodward, D. J., McIntire, D. D., Smith, W. K., … Mann, D. M. A. (1992). Disease-specific patterns of locus coeruleus cell loss. Annals of Neurology, 32(5), 667–676. https://doi.org/10.1002/ANA.410320510

Gray, S. R., Ye, L., Ye, J. Y., & Paukert, M. (2021). Noradrenergic terminal short-term potentiation enables modality-selective integration of sensory input and vigilance state. Science Advances, 7(51). https://doi.org/10.1126/SCIADV.ABK1378

Hammerschmidt, T., Kummer, M. P., Terwel, D., Martinez, A., Gorji, A., Pape, H. C., … Heneka, M. T. (2013). Selective loss of noradrenaline exacerbates early cognitive dysfunction and synaptic deficits in APP/PS1 mice. Biological Psychiatry, 73(5), 454–463. https://doi.org/10.1016/J.BIOPSYCH.2012.06.013

Heneka, M. T., Galea, E., Gavriluyk, V., Dumitrescu-Ozimek, L., Daeschner, J. A., O’Banion, M. K., … Feinstein, D. L. (2002). Noradrenergic depletion potentiates beta -amyloid-induced cortical inflammation: implications for Alzheimer’s disease. The Journal of Neuroscience : The Official Journal of the Society for Neuroscience, 22(7), 2434–2442. https://doi.org/10.1523/JNEUROSCI.22-07-02434.2002

Heneka, M. T., Ramanathan, M., Jacobs, A. H., Dumitrescu-Ozimek, L., Bilkei-Gorzo, A., Debeir, T., … Staufenbiel, M. (2006). Locus ceruleus degeneration promotes Alzheimer pathogenesis in amyloid precursor protein 23 transgenic mice. The Journal of Neuroscience : The Official Journal of the Society for Neuroscience, 26(5), 1343–1354. https://doi.org/10.1523/JNEUROSCI.4236-05.2006

Hirning, L. D., Fox, A. P., Mccleskey, E. W., Olivera, B. M., Thayer, S. A., Miller, R. J., & Tsien, R. W. (1988). Dominant Role of N-Type Ca2+ Channels in Evoked Release of Norepinephrine from Sympathetic Neurons. Science, 239(4835), 57–61. https://doi.org/10.1126/SCIENCE.2447647

Iwai, Y., Ozawa, K., Yahagi, K., Mishima, T., Akther, S., Vo, C. T., … Hirase, H. (2021). Transient Astrocytic Gq Signaling Underlies Remote Memory Enhancement. Frontiers in Neural Circuits, 15, 18. https://doi.org/10.3389/FNCIR.2021.658343/BIBTEX

Jackson, J. G., & Robinson, M. B. (2015). Reciprocal Regulation of Mitochondrial Dynamics and Calcium Signaling in Astrocyte Processes. Journal of Neuroscience, 35(45), 15199–15213. https://doi.org/10.1523/JNEUROSCI.2049-15.2015

Jankowsky, J. L., Fadale, D. J., Anderson, J., Xu, G. M., Gonzales, V., Jenkins, N. A., … Borchelt, D. R. (2004). Mutant presenilins specifically elevate the levels of the 42 residue beta-amyloid peptide in vivo: evidence for augmentation of a 42-specific gamma secretase. Human Molecular Genetics, 13(2), 159–170. https://doi.org/10.1093/HMG/DDH019

Jardanhazi-Kurutz, D., Kummer, M. P., Terwel, D., Vogel, K., Thiele, A., & Heneka, M. T. (2011). Distinct adrenergic system changes and neuroinflammation in response to induced locus ceruleus degeneration in APP/PS1 transgenic mice. Neuroscience, 176, 396–407. https://doi.org/10.1016/J.NEUROSCIENCE.2010.11.052

Kaeser, P. S., Deng, L., Wang, Y., Dulubova, I., Liu, X., Rizo, J., & Südhof, T. C. (2011). RIM proteins tether Ca2+ channels to presynaptic active zones via a direct PDZ-domain interaction. Cell, 144(2), 282–295. https://doi.org/10.1016/J.CELL.2010.12.029/ATTACHMENT/8C7BD067-CDB1-43BF-97AC-71BB4BDAE9F7/MMC1.PDF

Kalinin, S., Polak, P. E., Lin, S. X., Sakharkar, A. J., Pandey, S. C., & Feinstein, D. L. (2012). The noradrenaline precursor L-DOPS reduces pathology in a mouse model of Alzheimer’s disease. Neurobiology of Aging, 33(8), 1651–1663. https://doi.org/10.1016/J.NEUROBIOLAGING.2011.04.012

Kofuji, P., & Araque, A. (2021). Astrocytes and Behavior. Https://Doi.Org/10.1146/Annurev-Neuro-101920-112225, 44, 49–67. https://doi.org/10.1146/ANNUREV-NEURO-101920-112225

Kol, A., Adamsky, A., Groysman, M., Kreisel, T., London, M., & Goshen, I. (2020). Astrocytes contribute to remote memory formation by modulating hippocampal–cortical communication during learning. Nature Neuroscience 2020 23:10, 23(10), 1229–1239. https://doi.org/10.1038/s41593-020-0679-6

Krick, N., Ryglewski, S., Pichler, A., Bikbaev, A., Götz, T., Kobler, O., … Duch, C. (2021). Separation of presynaptic Cav2 and Cav1 channel function in synaptic vesicle exo- And endocytosis by the membrane anchored Ca2+ pump PMCA. Proceedings of the National Academy of Sciences of the United States of America, 118(28). https://doi.org/10.1073/PNAS.2106621118/-/DCSUPPLEMENTAL

Kuchibhotla, K. V., Lattarulo, C. R., Hyman, B. T., & Bacskai, B. J. (2009). Synchronous hyperactivity and intercellular calcium waves in astrocytes in Alzheimer mice. Science, 323(5918), 1211–1215. https://doi.org/10.1126/SCIENCE.1169096/SUPPL_FILE/KUCHIBHOTLA-SOM.PDF

Lee, Y., Messing, A., Su, M., & Brenner, M. (2008). GFAP promoter elements required for region-specific and astrocyte-specific expression. Glia, 56(5), 481–493. https://doi.org/10.1002/GLIA.20622

Lim, E. Y., Ye, L., & Paukert, M. (2021). Potential and Realized Impact of Astroglia Ca2 + Dynamics on Circuit Function and Behavior. Frontiers in Cellular Neuroscience, 15, 194. https://doi.org/10.3389/FNCEL.2021.682888/BIBTEX

Lockrow, J., Boger, H., Gerhardt, G., Aston-Jones, G., Bachman, D., & Granholm, A. C. (2011). A noradrenergic lesion exacerbates neurodegeneration in a Down syndrome mouse model. Journal of Alzheimer’s Disease : JAD, 23(3), 471–489. https://doi.org/10.3233/JAD-2010-101218

Louboutin, J. P., Chekmasova, A. A., Marusich, E., Chowdhury, J. R., & Strayer, D. S. (2010). Efficient CNS gene delivery by intravenous injection. Nature Methods 2010 7:11, 7(11), 905–907. https://doi.org/10.1038/nmeth.1518

Madisen, L., Garner, A. R., Shimaoka, D., Chuong, A. S., Klapoetke, N. C., Li, L., … Zeng, H. (2015). Transgenic mice for intersectional targeting of neural sensors and effectors with high specificity and performance. Neuron, 85(5), 942–958. https://doi.org/10.1016/J.NEURON.2015.02.022/ATTACHMENT/50FCEBAB-008B-4AE6-AE9D-46E1DE60A359/MMC5.MP4

Marcyniuk, B., Mann, D. M. A., & Yates, P. O. (1986). The topography of cell loss from locus caeruleus in Alzheimer’s disease. Journal of the Neurological Sciences, 76(2–3), 335–345. https://doi.org/10.1016/0022-510X(86)90179-6

Marcyniuk, B., Mann, D. M. A., & Yates, P. O. (1989). The topography of nerve cell loss from the locus caeruleus in elderly persons. Neurobiology of Aging, 10(1), 5–9. https://doi.org/10.1016/S0197-4580(89)80004-1

Marcyniuk, B., Mann, D. M. A., Yates, P. O., & Ravindra, C. R. (1988). Topography of nerve cell loss from the locus caeruleus in middle aged persons with Down’s syndrome. Journal of the Neurological Sciences, 83(1), 15–24. https://doi.org/10.1016/0022-510X(88)90016-0

Matthews, K. L., Chen, C. P. L. H., Esiri, M. M., Keene, J., Minger, S. L., & Francis, P. T. (2002). Noradrenergic changes, aggressive behavior, and cognition in patients with dementia. Biological Psychiatry, 51(5), 407–416. https://doi.org/10.1016/S0006-3223(01)01235-5

McCarty, D. M., DiRosario, J., Gulaid, K., Muenzer, J., & Fu, H. (2009). Mannitol-facilitated CNS entry of rAAV2 vector significantly delayed the neurological disease progression in MPS IIIB mice. Gene Therapy 2009 16:11, 16(11), 1340–1352. https://doi.org/10.1038/gt.2009.85

Mufson, E. J., Counts, S. E., Perez, S. E., & Ginsberg, S. D. (2008). Cholinergic system during the progression of Alzheimer’s disease: therapeutic implications. Expert Review of Neurotherapeutics, 8(11), 1703–1718. https://doi.org/10.1586/14737175.8.11.1703

Nagai, J., Bellafard, A., Qu, Z., Yu, X., Ollivier, M., Gangwani, M. R., … Khakh, B. S. (2021). Specific and behaviorally consequential astrocyte G q GPCR signaling attenuation in vivo with iβARK. Neuron, 109(14), 2256–2274.e9. https://doi.org/10.1016/J.NEURON.2021.05.023

Nagai, J., Rajbhandari, A. K., Gangwani, M. R., Hachisuka, A., Coppola, G., Masmanidis, S. C., … Khakh, B. S. (2019). Hyperactivity with Disrupted Attention by Activation of an Astrocyte Synaptogenic Cue. Cell, 177(5), 1280–1292.e20. https://doi.org/10.1016/J.CELL.2019.03.019/ATTACHMENT/43F2B535-2C38-4526-8664-079CAFF3C20D/MMC2.XLSX

Nimmerjahn, A., Mukamel, E. A., & Schnitzer, M. J. (2009). Motor behavior activates Bergmann glial networks. Neuron, 62(3), 400–412. https://doi.org/10.1016/J.NEURON.2009.03.019

Pabst, M., Braganza, O., Dannenberg, H., Hu, W., Pothmann, L., Rosen, J., … Beck, H. (2016). Astrocyte Intermediaries of Septal Cholinergic Modulation in the Hippocampus. Neuron, 90(4), 853–865. https://doi.org/10.1016/J.NEURON.2016.04.003

Papouin, T., Dunphy, J. M., Tolman, M., Dineley, K. T., & Haydon, P. G. (2017). Septal Cholinergic Neuromodulation Tunes the Astrocyte-Dependent Gating of Hippocampal NMDA Receptors to Wakefulness. Neuron, 94(4), 840–854.e7. https://doi.org/10.1016/J.NEURON.2017.04.021

Paukert, M., Agarwal, A., Cha, J., Doze, V. A., Kang, J. U., & Bergles, D. E. (2014). Norepinephrine controls astroglial responsiveness to local circuit activity. Neuron, 82(6), 1263–1270. https://doi.org/10.1016/J.NEURON.2014.04.038

Perez, S. E., Dar, S., Ikonomovic, M. D., DeKosky, S. T., & Mufson, E. J. (2007). Cholinergic forebrain degeneration in the APPswe/PS1ΔE9 transgenic mouse. Neurobiology of Disease, 28(1), 3–15. https://doi.org/10.1016/J.NBD.2007.06.015

Perry, E. K., Perry, R. H., Blessed, G., & Tomlinson, B. E. (1978). Changes in brain cholinesterases in senile dementia of Alzheimer type. Neuropathology and Applied Neurobiology, 4(4), 273–277. https://doi.org/10.1111/J.1365-2990.1978.TB00545.X

Pinto-Duarte, A., Roberts, A. J., Ouyang, K., & Sejnowski, T. J. (2019). Impairments in remote memory caused by the lack of Type 2 IP3 receptors. Glia, 67(10), 1976–1989. https://doi.org/10.1002/GLIA.23679

Polack, P.-O., Friedman, J., & Golshani, P. (2013). Cellular mechanisms of brain state-dependent gain modulation in visual cortex, 16(9). https://doi.org/10.1038/nn.3464

Reimer, J., Mcginley, M. J., Liu, Y., Rodenkirch, C., Wang, Q., Mccormick, D. A., & Tolias, A. S. (2016). Pupil fluctuations track rapid changes in adrenergic and cholinergic activity in cortex. https://doi.org/10.1038/ncomms13289

Rokosh, D. G., & Simpson, P. C. (2002). Knockout of the α1A/C-adrenergic receptor subtype: The α1A/C is expressed in resistance arteries and is required to maintain arterial blood pressure. Proceedings of the National Academy of Sciences of the United States of America, 99(14), 9474–9479. https://doi.org/10.1073/PNAS.132552699/SUPPL_FILE/5526SUPPINFO.PDF

Saija, A., Princi, P., De Pasquale, R., & Costa, G. (1989). Modifications of the permeability of the bloodbrain barrier and local cerebral metabolism in pentobarbital- and ketamine-anaesthetized rats. Neuropharmacology, 28(9), 997–1002. https://doi.org/10.1016/0028-3908(89)90202-5

Saito, T., Matsuba, Y., Mihira, N., Takano, J., Nilsson, P., Itohara, S., … Saido, T. C. (2014). Single App knock-in mouse models of Alzheimer’s disease. Nature Neuroscience 2014 17:5, 17(5), 661–663. https://doi.org/10.1038/nn.3697

Sakakibara, Y., Hirota, Y., Ibaraki, K., Takei, K., Chikamatsu, S., Tsubokawa, Y., … Iijima, K. M. (2021). Widespread Reduced Density of Noradrenergic Locus Coeruleus Axons in the App Knock-In Mouse Model of Amyloid-Amyloidosis. Journal of Alzheimer’s Disease, 82, 1513–1530. https://doi.org/10.3233/JAD-210385

Salehi, A., Faizi, M., Colas, D., Valletta, J., Laguna, J., Takimoto-Kimura, R., … Mobley, W. C. (2009). Restoration of norepinephrine-modulated contextual memory in a mouse model of Down syndrome. Science Translational Medicine, 1(7). https://doi.org/10.1126/SCITRANSLMED.3000258

Santello, M., Toni, N., & Volterra, A. (2019). Astrocyte function from information processing to cognition and cognitive impairment. Nature Neuroscience 2019 22:2, 22(2), 154–166. https://doi.org/10.1038/s41593-018-0325-8

Shekari, A., & Fahnestock, M. (2021). Cholinergic neurodegeneration in Alzheimer disease mouse models. Handbook of Clinical Neurology, 182, 191–209. https://doi.org/10.1016/B978-0-12-819973-2.00013-7

Shen, K. Z., & North, R. A. (1992). Muscarine increases cation conductance and decreases potassium conductance in rat locus coeruleus neurones. The Journal of Physiology, 455(1), 471–485. https://doi.org/10.1113/JPHYSIOL.1992.SP019312

Slezak, M., Kandler, S., Van Veldhoven, P. P., Van den Haute, C., Bonin, V., & Holt, M. G. (2019). Distinct Mechanisms for Visual and Motor-Related Astrocyte Responses in Mouse Visual Cortex. Current Biology : CB, 29(18), 3120–3127.e5. https://doi.org/10.1016/J.CUB.2019.07.078

Srinivasan, R., Huang, B. S., Venugopal, S., Johnston, A. D., Chai, H., Zeng, H., … Khakh, B. S. (2015). Ca(2+) signaling in astrocytes from Ip3r2(-/-) mice in brain slices and during startle responses in vivo. Nature Neuroscience, 18(5), 708–717. https://doi.org/10.1038/NN.4001

Srinivasan, R., Lu, T. Y., Chai, H., Xu, J., Huang, B. S., Golshani, P., … Khakh, B. S. (2016). New Transgenic Mouse Lines for Selectively Targeting Astrocytes and Studying Calcium Signals in Astrocyte Processes In Situ and In Vivo. Neuron, 92(6), 1181–1195. https://doi.org/10.1016/J.NEURON.2016.11.030

Stephen, T. L., Higgs, N. F., Sheehan, D. F., Awabdh, S. Al, López-Doménech, G., Arancibia-Carcamo, I. L., & Kittler, J. T. (2015). Miro1 Regulates Activity-Driven Positioning of Mitochondria within Astrocytic Processes Apposed to Synapses to Regulate Intracellular Calcium Signaling. Journal of Neuroscience, 35(48), 15996–16011. https://doi.org/10.1523/JNEUROSCI.2068-15.2015

Stobart, J. L., Ferrari, K. D., Barrett, M. J. P., Glück, C., Stobart, M. J., Zuend, M., & Weber, B. (2018). Cortical Circuit Activity Evokes Rapid Astrocyte Calcium Signals on a Similar Timescale to Neurons. Neuron, 98(4), 726–735.e4. https://doi.org/10.1016/J.NEURON.2018.03.050/ATTACHMENT/21AED263-C491-4705-8782-CC38E70D8FDF/MMC3.MP4

Sugihara, H., Chen, N., & Sur, M. (2016). Cell-specific modulation of plasticity and cortical state by cholinergic inputs to the visual cortex. Journal of Physiology-Paris, 110(1–2), 37–43. https://doi.org/10.1016/J.JPHYSPARIS.2016.11.004

Takano, T., Han, X., Deane, R., Zlokovic, B., & Nedergaard, M. (2007). Two-Photon Imaging of Astrocytic Ca2+ Signaling and the Microvasculature in Experimental Mice Models of Alzheimer’s Disease. Annals of the New York Academy of Sciences, 1097(1), 40–50. https://doi.org/10.1196/ANNALS.1379.004

Takata, N., Mishima, T., Hisatsune, C., Nagai, T., Ebisui, E., Mikoshiba, K., & Hirase, H. (2011). Astrocyte calcium signaling transforms cholinergic modulation to cortical plasticity in vivo. The Journal of Neuroscience : The Official Journal of the Society for Neuroscience, 31(49), 18155–18165. https://doi.org/10.1523/JNEUROSCI.5289-11.2011

Thal, S. C., Luh, C., Schaible, E. V., Timaru-Kast, R., Hedrich, J., Luhmann, H. J., … Zehendner, C. M. (2012). Volatile anesthetics influence blood-brain barrier integrity by modulation of tight junction protein expression in traumatic brain injury. PloS One, 7(12). https://doi.org/10.1371/J0URNAL.PONE.0050752

Thrane, A. S., Thrane, V. R., Zeppenfeld, D., Lou, N., Xu, Q., Nagelhus, E. A., & Nedergaard, M. (2012). General anesthesia selectively disrupts astrocyte calcium signaling in the awake mouse cortex. Proceedings of the National Academy of Sciences of the United States of America, 109(46), 18974–18979. https://doi.org/10.1073/PNAS.1209448109

Verkhratsky, A. (2019). Astroglial Calcium Signaling in Aging and Alzheimer’s Disease. Cold Spring Harbor Perspectives in Biology, 11(7). https://doi.org/10.1101/CSHPERSPECT.A035188

Williams, J. A., & Reiner, P. B. (1993). Noradrenaline hyperpolarizes identified rat mesopontine cholinergic neurons in vitro. Journal of Neuroscience, 13(9), 3878–3883. https://doi.org/10.1523/JNEUROSCI.13-09-03878.1993

Ye, L., Haroon, M. A., Salinas, A., & Paukert, M. (2017). Comparison of GCaMP3 and GCaMP6f for studying astrocyte Ca2+ dynamics in the awake mouse brain. PLOS ONE, 12(7), e0181113. https://doi.org/10.1371/JOURNAL.PONE.0181113

Ye, L., Orynbayev, M., Zhu, X., Lim, E. Y., Dereddi, R. R., Agarwal, A., … Paukert, M. (2020). Ethanol abolishes vigilance-dependent astroglia network activation in mice by inhibiting norepinephrine release. Nature Communications, 11(1). https://doi.org/10.1038/S41467-020-19475-5

Zorec, R., Parpura, V., & Verkhratsky, A. (2018). Preventing neurodegeneration by adrenergic astroglial excitation. The FEBS Journal, 285(19), 3645–3656. https://doi.org/10.1111/FEBS.14456

