## Supplementary_Material for "Impaired neuromodulator crosstalk delays vigilance-dependent astroglia Ca^2+^ activation in mouse models of Alzheimer’s disease"

### Supplementary Figures

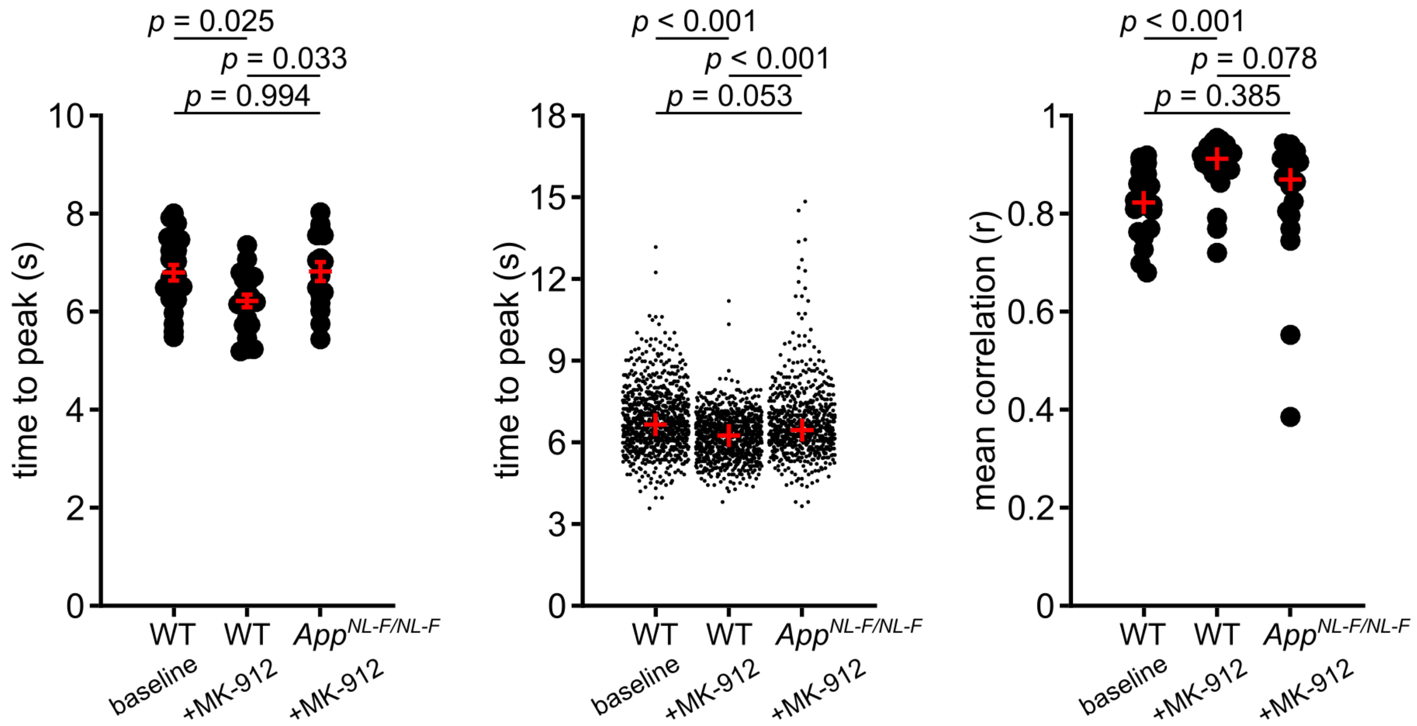


**Supplementary Figure 1 Enhancing norepinephrine release by blocking $\alpha_2$-adrenergic receptors with MK-912 partially rescued *App*$^{NL-F}$ phenotype in vigilance-dependent astrocyte $Ca^{2+}$ dynamics.** Population data: **Left**, time to peak in aged wildtype mice, *n* = 22 FOVs, 8 mice; and in aged *App*$^{NL-F/NL-F}$ mice, *n* = 16 FOVs, 7 mice; one-way ANOVA followed by Tukey-Kramer correction ($F(2,57) = 4.779$, $p = 0.012$). **Center**, time to peak of individual ROIs in aged wildtype mice, *n* = 824 ROIs, 8 mice; and in aged *App*$^{NL-F/NL-F}$ mice, *n* = 678 ROIs, 7 mice; Kruskal-Wallis test followed by Tukey-Kramer correction ($p < 0.001$). **Right**, mean correlation coefficient across ROIs in aged wildtype mice, *n* = 23 FOVs, 8 mice; and in aged *App*$^{NL-F/NL-F}$ mice, *n* = 18 FOVs, 7 mice; Kruskal-Wallis test followed by Tukey-Kramer correction ($p < 0.001$). If all data follow Gaussian distribution red symbols indicate mean ± SEM, if not red symbols represent median.

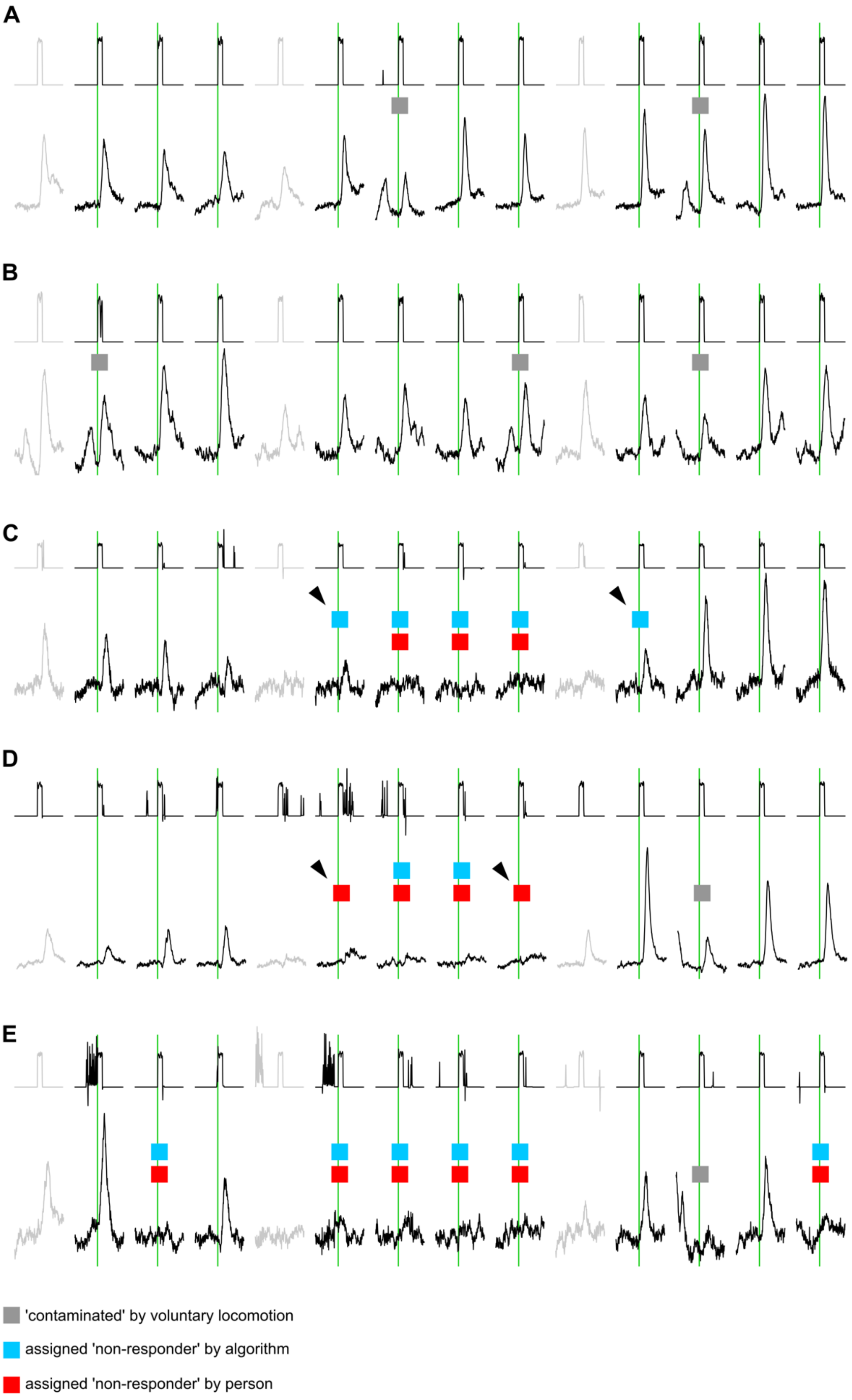


**Supplementary Figure 2 Criteria for eliminating trials of *App*$^{NL\text{-}F/NL\text{-}F}$ mice and wildtype littermates for analysis of time to peak.** (**A-B**) Two representative $Ca^{2+}$ traces showing repetitions indicated with grey boxes that were contaminated by voluntary locomotion prior to treadmill activation (green bars) and the start of enforced locomotion. (**C**) A representative $Ca^{2+}$ trace showing two repetitions (black arrows) where the automated algorithm assigned a non-responding event that was deemed responsive by human selection. (**D**) A representative $Ca^{2+}$ trace showing two repetitions (black arrows) where the automated algorithm failed to detect what people considered non-responding events. (**E**) A representative $Ca^{2+}$ trace showing correct assignment by the algorithm.

### Supplementary Methods

#### Experimental design

All animal procedures were conducted in accordance with guidelines and protocols of the University of Texas Health Science Center at San Antonio (UTHSCSA) Institutional Animal Care and Use Committee. At the time of data collection, aged APP/PS1(+) mice and wildtype littermates were between 17-22 months old, *App*$^{NL\text{-}F/NL\text{-}F}$ and age-matched wildtype mice were between 16-22 months old, and all other mice were between 2 and 7 months old. For all *in vivo* 2P experiments, we used a previously reported paradigm of head-restrained mouse on motorized treadmill (Paukert et al., 2014). $Ca^{2+}$ dynamics in astrocytes of left V1 were observed during bouts of enforced locomotion lasting either 2 minutes (Figures 1-4) or 5 seconds (Figure 5 and Supplementary Figures 1 and 2) at 90-110 mm/s. Locomotion protocols and data analysis routines described below were automated to minimize risk of experimenter bias. In global knockout experiments, the experimenter was uninformed of mouse genotype. On the basis of availability, male and female mice were assigned randomly to individual experiments, and most datasets contain data from mice of both sexes, except for APP/PS1(+) mice where only females were available. Sex-disaggregated data analysis revealed no trends attributable to sex. For most experiments (Figures 1-4), one FOV was imaged per mouse and statistical analysis was based on number of mice. For *App*$^{NL\text{-}F/NL\text{-}F}$ experiments (Figure 5 and Supplementary Figures 1 and 2), 1-3 FOVs were imaged per mouse and statistical analysis was based on number of FOVs.

#### Animals

Mice were kept in the Laboratory Animal Resources facility with ambient 72–78 °F temperature and 30–70% humidity and had ad libitum access to water and chow. Mice were maintained on a reverse 12h-light/12h-dark schedule (lights off at 9 a.m., on at 9 p.m.), and all experimental procedures were completed during the dark cycle. For *App*$^{NL\text{-}F/NL\text{-}F}$ mice and age-matched wildtype mice (Figure 5 and Supplementary Figures 1 and 2), GCaMP3 was virally expressed in astrocytes (see Virus paragraph) as genetically-encoded $Ca^{2+}$ indicator (GECI). Wildtype mice used as controls for *App*$^{NL\text{-}F/NL\text{-}F}$ mice were C57BL/6J and age-matched to the *App*$^{NL\text{-}F/NL\text{-}F}$ mice, which had been maintained on C57BL/6J background. For all other experiments, GCaMP6f (Ai95) (Madisen et al., 2015) or Lck-GCaMP6f (Srinivasan et al., 2016) was expressed as GECI in a Cre-dependent manner. Each animal was heterozygous for one GECI allele and heterozygous for one Cre recombinase allele expressed specifically in astroglia (Figures 1-3; *Slc1a3-CreER*$^{T}$ (Paukert et al., 2014)) or in noradrenergic neurons (Figure 4; *Dbh-Cre* (Gerfen et al., 2013)). For global *Adra1a* (Rokosh & Simpson, 2002) knockout experiments, the final breeding step produced offspring heterozygous for GCaMP6f, heterozygous for *Slc1a3-CreER*$^{T}$, and either homozygous or heterozygous for or lacked the wildtype *Adra1a* gene. *Adra1a* mice were born in a frequency expected of Mendelian principles. All genotypes were determined from extracted toe or tail sample DNA via polymerase chain reaction (PCR) using gene-specific primers.

##### Tamoxifen administration and recombination efficiency

Tamoxifen (Sigma-Aldrich, #T5648-5G) was freshly dissolved in sunflower seed oil (Sigma-Aldrich, #S5007) at a concentration of 10 mg/ml by flicking and sonication for approximately 30 min and was stored at 4˚C for up to 5 days. For experiments using the *Slc1a3-CreER*$^{T}$ mouse line overexpressing GCaMP6f (Figures 1-3), tamoxifen was injected i.p. (100 µL per 10 g mouse body weight starting at age 3–4 weeks) three times within 5 days. For experiments using the *Dbh-Cre* mouse line to overexpress Lck-GCaMP6f in noradrenergic neurons (Figure 4), no tamoxifen administration was necessary and from immunofluorescence analysis of LC, recombination efficiency is close to 100% (Ye et al., 2020). Surgeries were performed 1–2 weeks after the last tamoxifen injection, and experiments started at least 2 weeks following surgeries.

##### Animal surgery

Surgical installation of chronic cranial windows was comprised of two steps. In the first step, mice were placed under anesthesia with 100 mg/kg ketamine and 10 mg/kg xylazine administered i.p. and positioned on a heating pad to maintain body temperature at ~36 °C. Hair was removed and then skin disinfected using povidone-iodine. Skin and muscles were removed from the skull, and 3% hydrogen peroxide was applied to disinfect and prevent bleeding. The periosteum was shaved off, and remaining muscle surrounding the exposed skull was covered with a thin layer of cyanoacrylate cement. A custom-designed stainless steel head-plate with a 4 mm x 6 mm oval opening was centered above left primary visual cortex at lambda, 2.5 mm lateral from midline and mounted on the skull using dental cement (C&B Metabond, Parkell Inc., Brentwood). Wound edges were coated with Neosporin® ointment containing pramoxine, and mice were placed in a heated cage during recovery. In the second-step surgery 3-7 days later, under isoflurane anesthesia (1.5–3% vol./vol. isoflurane in $O_2$ with flow rate adjusted according to hindpaw pinch reflex) and on a heating pad, mice underwent a craniotomy where a 2 mm x 2 mm portion of skull within the oval opening of the steel head-plate was removed. Dura mater was torn and pushed aside, and the exposed brain tissue was covered with a glass window comprised of three layers of No. 1 Corning cover glass fused to each other by UV-curable Norland Optical Adhesive #81. The window edges were sealed to the skull with dental cement (Ortho-Jet-Acrylic-Powder, Lang) with small amount of pooled CSF preventing direct contact of dental cement with brain tissue. After 1 week of post-surgery recovery, mice were habituated to the linear treadmill and recording conditions for 3 sessions, exactly replicating the imaging experiment except that saline was injected i.p. instead of drugs and the laser was kept off. Imaging was performed at least 2 weeks after surgeries.

**Virus**

To express GCaMP in astrocytes of *App*$^{NL\text{-}F/NL\text{-}F}$ mice and age-matched wildtype mice (Figure 5 and Supplementary Figures 1 and 2), GCaMP3 was virally expressed specifically in astrocytes. Comparison of GCaMP3 and GCaMP6f reveal only minor differences in sensor properties and that either variant is suitable for studying *in vivo* astroglia $Ca^{2+}$ dynamics (Ye et al., 2017). The plasmid was originally made by replacing the coding region for mKate2.5f in the AAV-GFAP-mKate2.5f plasmid (a gift from V. Gradinaru, California Institute of Technology, Pasadena, CA, now available from Addgene 99129) with the coding region for GCaMP3 by PCR-amplifying this region out of the Zac2.1 gfaABC1D-Cyto-GCaMP3 plasmid (a gift from B. Khakh; Addgene, 44331) and digesting and ligating the two fragments. AAV-gfaABC1D-GCaMP3 was packaged in the AAV-PHP.B capsid (a gift from V. Gradinaru, now available from Addgene, 103002), which is optimized for CNS transduction (Deverman et al., 2016). Virus packaging was performed by Penn Vector Core in the Gene Therapy Program of the University of Pennsylvania who provided 900 µL at $2.77 \times 10^{13}$

genome copies per mL (GC/mL, detected by a droplet digital PCR-based method). Shortly before injection, stock virus solution was diluted in D-PBS 20-fold to a working concentration of $1.385 \times 10^{12}$ GC/mL. To improve blood-brain-barrier permeability, D-mannitol (VWR cat no. 0122) was administered by i.p. injection (25% w/v in 0.9% saline at 30 μL/g body weight) 20 minutes before virus administration (Louboutin, Chekmasova, Marusich, Chowdhury, & Strayer, 2010; McCarty, DiRosario, Gulaid, Muenzer, & Fu, 2009), and isoflurane anesthesia rather than ketamine/xylazine was used (Saija, Princi, De Pasquale, & Costa, 1989; Thal et al., 2012). 50 μL of virus was retro-orbitally injected free-hand with a 30G, 25 mm needle through the medial canthus into the retro-orbital sinus of the left eye. Mice were habituated 3 weeks later and imaged 4 weeks after retro-orbital injection of virus.

##### *In vivo* 2P imaging

Two setups were used for $Ca^{2+}$ imaging experiments. For APP/PS1 *in vivo* experiments (Figure 1), a galvanometer-based 2P laser-scanning Movable Objective Microscope (MOM) (Sutter Instruments) with a 16x, 0.80 NA water-immersion objective (Nikon) was used. A pulsed Ti:Sapphire laser beam at 920 nm, 80 MHz repetition rate, 140 fs pulse width (Coherent Inc., Chameleon Ultra II) was focused at approximately 60 μm below the pial surface in layer 1 of primary visual cortex. To minimize brain injury, laser power was attenuated to 10–35 mW at the front aperture of the objective. Emitted light was detected using a gallium arsenide phosphide (GaAsP) photomultiplier tube (H10770PA-40; Hamamatsu Photonics). The awake mouse was placed on a custom-made linear treadmill, and the head-plate was secured under the microscope objective. The speed of the treadmill belt was monitored with an optical encoder. The belt was freely movable so that the mouse could walk voluntarily, and also at predefined episodes, a servo motor could be engaged to enforce locomotion at 90–110 mm/s. Electromyography (EMG) signals were recorded as the surface potential difference between two silver wires inserted subcutaneously at the right shoulder and left hip of the mouse. The 2P microscope was controlled by an Xi Computer Corporation personal computer (Intel(R) Core(TM) i7-4930 CPU @ 3.40 GHz, 8 GB of RAM) running ScanImage (v3.8.1; Vidrio Technologies, LLC) software within MATLAB R2011b (Mathworks). Image acquisition was triggered at a rate of 1.5 frames/s, and locomotion speed data, EMG data, and Y-mirror position data were simultaneously acquired at 20 kHz sampling rate using National Instruments boards controlled by custom-written scripts in

Labview2013 (version 13.0.1f2, National Instruments). Acquired frames were 400 µm x 400 µm at a resolution of 512 pixels per line and 512 lines per frame. Non-imaging data were post hoc downsampled to the image acquisition frame rate and the Y-mirror signal was used to assign appropriate data bins to individual image frames. The entire experimental setup was enclosed in a blackout box.

For all other imaging experiments (Figures 2-5 and Supplementary Figures 1 and 2), a resonant scanning version of the MOM (Sutter Instruments) was used with a 16x, 0.80 NA water-immersion objective (Nikon). A pulsed Ti:Sapphire laser beam at 920 nm, 80 MHz repetition rate, <120 fs pulse width (Insight DS+, Spectra-Physics MKS Instruments Light & Motion) was used for 2P excitation. Data were acquired using volumetric scanning with a step size of 10 µm, 6 slices per volume covering 50 µm in the z-plane, and at 30 frames per second and 4.29 volumes per second. Acquired frames were 400 µm x 400 µm at a resolution of 512 pixels per line and 512 lines per frame. Laser power was adjusted to 10–35 mW at the front aperture of the objective. The microscope was controlled by an Xi Computer Corporation personal computer (Intel(R) Core(TM) i7-5930K CPU @ 3.50 GHz, 16 GB of RAM) running ScanImage (v5.0; Vidrio Technologies, LLC) software within MATLAB R2016a (Mathworks). All other design principles, including that of the treadmill, were identical between microscopes.

**Pharmacology**

All drugs were applied systemically via i.p. injection. The following drug preparations were used and administered at 100 µl/10 g body weight. For a target dosage of 3 mg/kg scopolamine, stock scopolamine solution was made at 0.25 mg/mL by mixing 1 mg in 4 mL of 0.9% saline, and 6 µL of stock was mixed with 494 µL saline. For a target dosage of 0.03 mg/kg MK-912, 3 mg of MK-912 was dissolved in 1 ml of saline and then diluted 1000x in saline. For control i.p. injections, 100 µl saline/10 g body weight was injected. Injections were conducted immediately following the previous imaging trial. The same FOV was imaged before and after i.p. injections. Two versions of scopolamine and two versions of MK-912 were used due to discontinued production of the original versions. (-)-scopolamine hydrochloride (Sigma S1013) and (-)-scopolamine hydrobromide trihydrate (Sigma S1875) were used. MK-912 hydrochloride hydrate (Sigma M7065, was temporarily discontinued and currently available again) and MK-912 hydrate (Santa Cruz Biotechnology sc-253050) were used.

**Data analysis**

Imaging data were saved in ScanImage as tiff files, imported to MATLAB R2016a or R2018b, and stored as mat files. Analysis was conducted using custom-written scripts employing a combination of built-in, open-source, and custom-written functions. Any computer code can be shared upon request. For experiments acquired with volume scanning (Figures 2-5 and Supplementary Figures 1 and 2), each volume was compressed into a single maximum intensity projection and analyzed as a single frame series of 4.29 frames per second. Images were first passed through a Gaussian filter (1.52 SD per pixel distance) to attenuate random noise of the detector. Individual frames within the entire time series (including all imaging trials with each trial representing: baseline imaging frames + imaging frames during 5 or 120 s locomotion episode + imaging frames until the next imaging pause highlighted by black bars in all pseudocoloured plots) were registered to maximize correlation. For noradrenergic terminals, where individual structure elements could not be assigned to individual cells, ROIs were spatially defined by an 8 x 8 checkerboard pattern, which resulted in ROIs of approximately 50 μm x 50 μm (Figure 4). For astrocyte experiments (Figures 1, 2, 3, 5, and Supplementary Figures 1 and 2), ROIs were drawn around individual astrocytes encompassing the soma and fine processes, and then a custom, automated script detected the dimmest pixels that failed to reach a minimum threshold of fluorescence and removed them from within the drawn ROIs. For each ROI, a mean raw fluorescence value of all pixels was corrected by subtracting the average detector offset obtained from imaging under identical conditions with the laser shutter closed. For each ROI and frame this corrected fluorescence value (F) was normalized to baseline by calculating $(F - F_{median})/F_{median}$ where $F_{median}$ was the median fluorescence value of the respective ROI during baseline (from start of a trial until the frame before onset of locomotion). For all analyzed parameters individual ROIs were analyzed first and if not reported individually (Figure 5 F and G) $\Delta F/F$ values from all ROIs of a FOV were averaged. The term "$Ca^{2+}$ change" represents mean $\Delta F/F$ within 20 s from onset of locomotion. "Time to peak" (TTP) represents the time from onset of locomotion to the maximum $\Delta F/F$ value within 20 s from onset of locomotion. The "mean correlation coefficient" represents the mean Pearson's linear correlation coefficient among all pairs of ROI $\Delta F/F$ traces within 20 s from onset of locomotion. We previously found that voluntary, locomotion-induced Bergmann glia $Ca^{2+}$ elevations during the baseline phase can suppress enforced locomotion-induced responses (Paukert et al., 2014). Therefore, trials with episodes of voluntary locomotion preceding treadmill activation were eliminated from analysis, and these trials were identified as those where at least two consecutive $\Delta F/F$ values within baseline phase exceeded 3x the standard deviation (SD) of baseline $\Delta F/F$ values averaged across all trials of a dataset. For each experiment, values from trials that were not contaminated by voluntary activity preceding treadmill activation were averaged and represented as dot in the summary plots. For all experiments, the first repetition was eliminated from analysis, and also the first repetition following i.p. injection of a drug was eliminated to account for the time required for the drug to reach its effective concentration in the brain. For experiments that used viral delivery of GCaMP3 (Figure 5 and Supplementary Figures 1 and 2), we occasionally noticed in individual trials that the astrocyte population of a FOV failed to respond (out of 382 non-contaminated trials, 21 showed no response). Since time to peak could not be determined in these cases, they were excluded for analysis of this parameter. We detected 'non-responding' trials by combining an automated algorithm with independent human judgement. First, the custom script used the same strategy described above for detecting trials contaminated by voluntary activity-induced $Ca^{2+}$ elevations and scanned for such events during the entire repetition; if no event was detected at any point during the repetition, this was assigned as 'non-responding.' In parallel, two colleagues in the lab who were blind to genotype were given a random assortment of $Ca^{2+}$ traces and selected trials that they deemed 'non-responding.' The automated results were assessed post-hoc compared to the assignments by people. In total, 21 trials were deemed non-responsive; of these, 5 were undetected by the algorithm but were assigned by people, and 6 repetitions that were flagged by the algorithm were considered 'responsive' and included in the analysis (Supplementary Figure 2). If a response was detectable, the time to peak was analyzed.

**Statistical analysis**

Statistical analyses were performed using MATLAB R2018b (MathWorks). For each group of a data set the Lilliefors test was applied to test for Gaussian distribution. If all groups followed a Gaussian distribution, the data was presented with mean ± SEM. If two unrelated groups were compared, the two-tailed Student's *t*-test was used; if two groups were related, the two-tailed paired Student's *t*-test was used. For more than two unrelated groups, one-way ANOVA was used; and if groups were related, repeated-measures ANOVA was used. If at least one group did not follow Gaussian distribution the data was presented with median. If two or more unrelated groups were compared, the Kruskal Wallis test was used; if more than two groups were compared and were related, the Friedman test was applied. For all tests that involved more than one comparison, Tukey–Kramer correction was applied for multiple comparisons. The sample number for individual tests and respective test type applied are mentioned in the results section and also in the respective figure legends. Test results are indicated by *p* values and a significance level of 0.05 was applied.
